# Cross-modal Denoising and Integration of Spatial Multi-omics data with CANDIES

**DOI:** 10.1101/2025.04.17.649333

**Authors:** Ye Liu, Wanpeng Zou, Yuekai Li, Jiayi Wang, Mingxuan Cai, Hongmin Cai

**Author notes:** Contributing authors.

## Abstract

Spatial multi-omics data offer a powerful framework for integrating diverse molecular profiles while maintaining the spatial organization of cells. However, inherent variations in data quality and noise levels across different modalities pose significant challenges to accurate integration and analyses. In this paper, we introduce CANDIES, which leverages a conditional diffusion model and contrastive learning to effectively denoise and integrates spatial multi-omics data. With our innovative model and algorithm designs, CANDIES not only enhances the quality of spatial multi-omics data, but also yields a unified and comprehensive joint representation, thereby empowering many downstream analysis. We conduct extensive evaluations on diverse synthetic and real datasets, including spatial CITE-seq data from human skin biopsy tissue, MISAR-seq data from the mouse brain, spatial ATAC-RNA-seq data from the mouse embryo and 10× visium data from human lymph nodes. CANDIES shows superior performance on various downstream tasks, including denoising, spatial domain identification, spatiotemporal trajectories reconstruction, and spatial association mapping for complex human traits. In particular, we show that CANDIES representations can be integrated with the rich resources from genome-wide association studies (GWASs), allowing the spatial domains to be linked with complex human traits, yielding spatially resolved interpretation of complex traits in their relevant tissues.

## Introduction

Groundbreaking spatially resolved omics technologies have become powerful tools for understanding cellular states and cellular organization within tissue. Recent advancements in spatial multi-omics techniques have enabled the simultaneous spatial measurement of multiple molecular profiles within the same tissue section [1]. For example, co-detection methods for protein and gene expression include DBiT-seq [2], Spatial CITE-seq [3], Stereo-CITE-seq [4], and CosMxTM Spatial Molecular Imager (SMI) [5]. Additionally, the comapping of the chromatin accessibility and gene expression is achieved by spatial ATAC–RNA-seq [6] and MISAR-seq [7]. This development allows for a more comprehensive understanding of the molecular landscape within tissues, offering new insights into tissue organization, function, and disease mechanisms [8].

Despite advancements in spatial multi-omics technologies, analyzing spatial multi-omics data remains highly challenging. One major hurdle is the inherent noise within each modality, which can obscure meaningful biological signals and hinder accurate data integration. For instance, technical noise introduced during spatial transcriptomic sequencing, coupled with false zero counts in gene expression measurements due to low RNA capture efficiency, poses major obstacles to accurate analysis and downstream applications [9, 10, 11]. Second, extracting data from one modality might disrupt the molecular stability required for another, which could diminish the accuracy of subsequent molecular sequencing and lead to inconsistencies in data quality across different modalities [12]. For example, due to suboptimal antibody-derived tags (ADTs) concentrations and enzymatic reaction parameters, competition between ADTs and mRNAs during reverse transcription can reduce transcript detection efficiency compared to single-modality spatial transcriptomics [3]. Third, each modality provides unique insights into cellular identity, offering complementary yet distinct advantages and limitations. The integration of multi-omics data is needed to leverage these diverse perspectives to improve the precision and sensitivity of identifying cell types and states, ultimately enabling a more comprehensive understanding of cellular heterogeneity and function [13].

Several methods, such as MultiVI [14], totalVI [15], and scCross [16] have been developed to integrate single-cell multi-omics data through deep generative models. For instance, scCross employed a bidirectional aligner and a discriminator for high-quality modality generation, allowing researchers to derive data from one modality based on another. However, biological molecular regulatory relationships are not solely dependent on molecular characteristics, they are also influenced by the tissue microenvironment [17]. Recently, many computational algorithms, including SpaGCN [18], GraphST [19], SEDR [20] and Giotto [21], have been developed for spatially resolved data. But these tools are designed for specific data modalities, such as spatial transcriptomics (ST) [22, 23] or spatial proteomics [24], often providing a fragmented view of the cellular landscape. Although these methods can be extended to spatial multi-omics data by concatenating features from various modalities, they remain suboptimal due to the inherent heterogeneity across these modalities. The field of spatial multi-omics integration is still in its early stages, with several recent methods such as SpatialGlue [25], PRESENT [26], COSMOS [27] and PRAGA [28], attempting to address the challenges of cross-modality representation with spatial context. However, these methods are often sensitive to noise, which can degrade their performance and limit their ability to uncover biologically meaningful insights.

To overcome these limitations, we proposed CANDIES, a novel framework leveraging <u>C</u>onditional diffusion model <u>A</u>nd co<u>N</u>trastive learning, specifically designed for cross-modal <u>D</u>enoising and <u>I</u>nt<u>E</u>gration of <u>S</u>patial multiomics data. By taking spatial multi-omics data with modalities exhibiting varying levels of noise as input, CANDIES refines the lower-quality modality by leveraging information from the higher-quality modality and spatial context. It then integrates spatial and enhanced feature information across modalities to produce robust, unified representations of spatial multi-omics data. CANDIES significantly improves the accuracy and reliability of downstream analysis, including denoising, spatial domain identification, pseudo-Spatiotemporal Map (pSM) construction, and spatially resolved mapping of complex human traits. To validate its effectiveness, we first evaluated CANDIES on a simulated dataset and spatial transcriptome-protein data from human skin biopsy tissue, where it outperformed state-of-the-art methods in both spatial single-modality denoising and multi-modal integration across a wide range of noise levels. Further analysis on spatial epigenome-transcriptome data from mouse brain and embryo demonstrated that CANDIES not only accurately distinguished major biological categories but also preserved the integrity of rare cell types. Lastly but importantly, CANDIES output can be integrated with summary statistics of genome-wide association studies (GWASs) to identify spatially resolved tissue domains associated with complex human traits. We integrated CANDIES results from spatial epigenome-transcriptome data from mouse brain and embryo, as well as spatial transcriptome-protein data from human lymph nodes with GWASs from 32 complex traits, uncovering interpretable associations between traits and relevant tissue regions. These results highlight CANDIES’ superior capability to extract biologically meaningful insights, providing a powerful framework for advancing the understanding of spatially resolved molecular landscapes.

## Results

### CANDIES model overview

CANDIES, shown in Fig. 1, is a comprehensive and innovative framework specifically designed for spatial multi-omics data with modalities exhibiting different levels of noise. As a two-stage method, CANDIES first refines the lower-quality modality by leveraging its spatially resolved dependence with the higher-quality modality. Subsequently, CANDIES facilitates more precise and robust integration of enhanced spatial multi-omics data by cross-modal alignment and multi-relation fusion, enabling a unified and biologically meaningful representation. By combining these stages, CANDIES effectively mitigates noise and improves the overall effectiveness of spatial multi-omics data analysis.

**Fig. 1:**
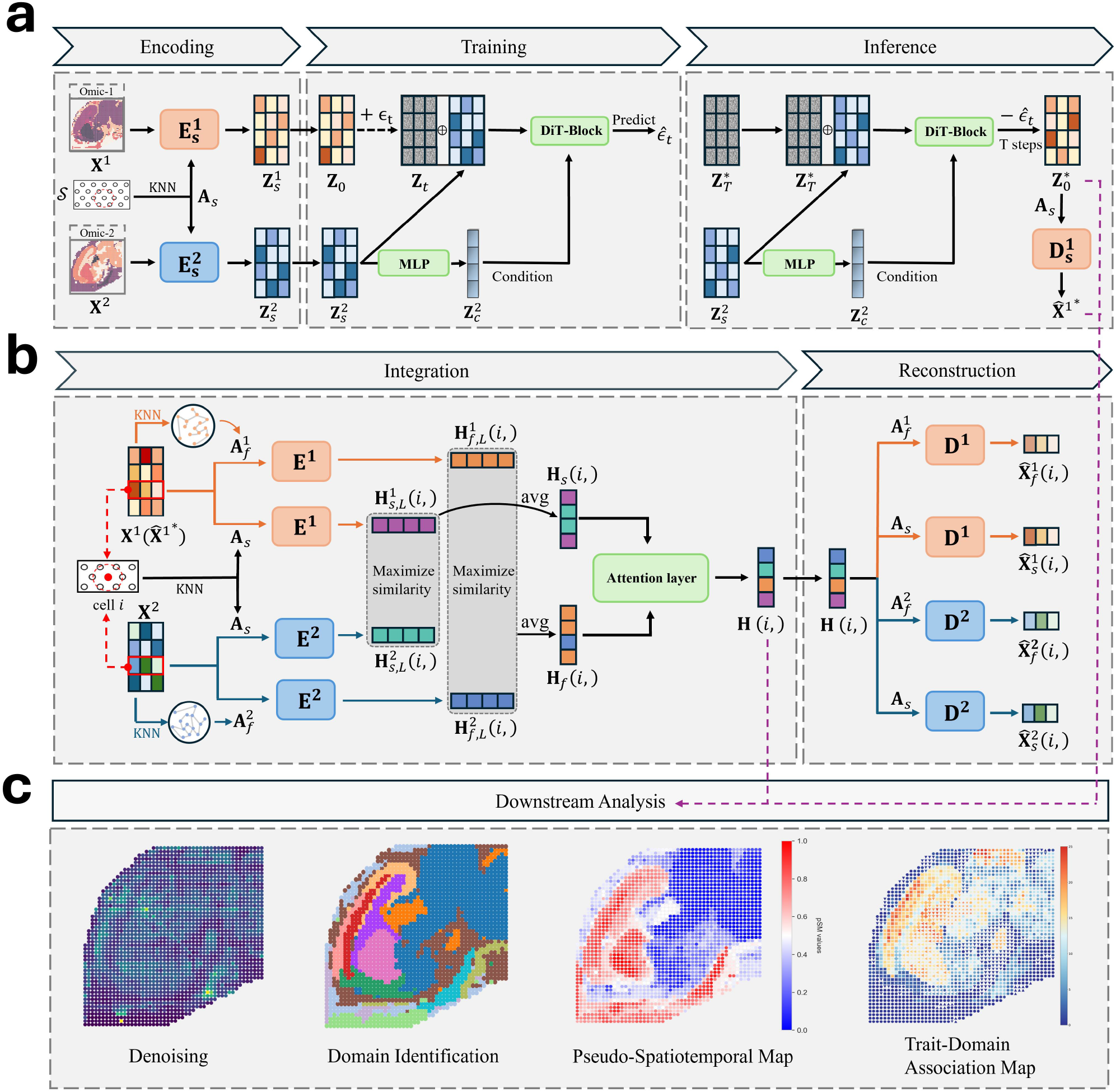
The framework of CANDIES model. **a**, The denoising phase of CANDIES. CANDIES begins by constructing a spatial graph using the k-nearest-neighbor (KNN) algorithm based on spatial coordinates. A graph auto-encoder (GAE) is then applied to this spatial graph with the features of each modality to generate low-dimensional embeddings for each modality, which facilitate the efficient training of the denoising model. Next, a diffusion transformer (DiT)-based conditional diffusion model enhances the embeddings of the lower-quality modality by leveraging its spatial dependencies with the higher-quality modality embeddings. Finally, the denoised embeddings of the lower-quality modality are decoded back into the original data space. **b**, The integration phase of CANDIES. CANDIES encodes denoised spatial multi-omics data into latent space by applying a dual graph convolution network (GCN) on both spatial proximity and feature similarity graphs for each omic modality. Subsequently, CANDIES aligns cross-modality embeddings via contrastive learning, and dynamically integrate spatial graph and feature graph through the attention mechanism, effectively preserving cellular heterogeneity while maintaining an optimal balance between spatial organization and molecular feature information. **c**, CANDIES supports four downstream analysis including denoising, spatial domain identification, pseudo-Spatiotemporal Map (pSM), and spatially resolved mapping of complex human traits.

In the first stage (Fig. 1a), CANDIES enhances the lower-quality modality data by a diffusion transformer (DiT)-based conditional diffusion model. Specifically, CANDIES first constructs a spatial graph based on the shared spatial coordinates, then a dual Graph Auto-Encoder (GAE) is trained to extract complementary information from the spatial graph with features of multiple modalities, generating embeddings for all spots within each modality. In the proposed DiT-based conditional diffusion model, the embeddings of the lower-quality modality are initialized as the starting point of the diffusion process, while the embeddings of the higher-quality modality serve as conditions to guide the reverse process. During the reverse denoising process, the embeddings of the lower-quality modality and the higher-quality modality are concatenated and fed into a DiT architecture [29], which acts as the backbone to capture long-range interactions and intricate patterns among spots. To enhance the conditioning effect, the embeddings of the higher-quality modality are further processed through a Multi-Layer Perceptron (MLP), generating refined representations to serve as the conditional input. This design combines the systematic refinement of noisy modality and the preservation of spatial dependencies, ensuring that the reliable information contained in the higher-quality modality is effectively leveraged to guide the denoising of the lower-quality modality, thereby resulting in denoised embeddings that are accurate and biologically interpretable. Finally, the pre-trained decoder maps the enhanced embeddings of the lower-quality modality back to the original feature space, enabling effective reconstruction of the lower-quality modality data.

In the integration stage (Fig. 1b), CANDIES constructs a joint low-dimensional representation by taking the enhanced spatial multi-omics data as input. To capture the shared biological signals among features, we construct a feature similarity graph for each modality with KNN. A dual Graph Convolutional Networks (GCN) [30] is then employed to integrate information from the feature graph and the spatial graph, producing latent embeddings for each spot within each modality. Unlike existing methods that directly work with the original omics data, CANDIES operates on the refined omics dataset with reduced noise, allowing the feature proximity to be better represented. To optimally align heterogeneous multi-modal information, contrastive learning is leveraged to encourage similarity of spatial and feature embeddings between the spots of different modalities while allowing heterogeneity between non-matching spots. This ensures that shared biological signals are captured while preserving modality-specific variations. Furthermore, an attention mechanism is introduced to integrate spatial and feature information by dynamically adjusting the contributions of the spatial and feature graphs. Such flexibility enables the model to adaptively determines the relative importance of spatial coherence and feature-level similarity. Finally, CANDIES uses spatial and feature graphs to reconstruct the expression profiles, enforcing the learned latent representations to preserve original expression patterns.

### CANDIES enhances spatial domain segmentation via denoising on simulated spatial multi-omics data

We conducted a comprehensive benchmarking study to evaluate the performance of CANDIES against several state-of-the-art approaches using simulated data. Following SpatialGlue [25], we generated an RNA modality with lower quality and a protein modality with higher quality (see details in Methods section). We controlled the quality of RNA modality by adjusting the dropout rate and the standard deviation of Gaussian noise added to the true expression level. Specifically, we varied the dropout rate for the RNA modality within [0.2, 0.5]. The Gaussian noise had a mean of 2.0 for both modalities, with the standard deviation varied within [2.0, 5.0] for RNA modality and fixed at 1.0 for protein modality (detailed parameters in the Supplementary Tables. S3-S4).We first demonstrated the capability of CANDIES to denoise the RNA modality by performing spatial domain identification based on the latent embeddings obtained from the refined RNA modality in the first step of CANDIES. We compared our method with various representative ST analysis methods, including GAAEST [31], GraphST [19], SEDR [20], SCAN-IT [32], SpaceFlow [33], SpaGCN [18], and STAGATE [34]. The performance was assessed using common clustering evaluation metrics, including Mutual Information (Mutual Info), Normalized Mutual Information (NMI), Adjusted Mutual Information (AMI), V-measure, Homogeneity, Completeness, Adjusted Rand Index (ARI) and Fowlkes-Mallows Index (FMI).

Fig. 2b shows the results obtained by setting the noise standard deviation as 3.0 and the dropout rate as 0.3 for ST data. As we can observe, CANDIES successfully recovered all four spatial factors of ST data with an ARI of 0.92, which is 3.7 times higher than the ARI of original ST data. Furthermore, while SEDR is also designed for ST data denoising, it cannot leverage the higher-quality protein modality to refine the RNA modality, we observed that CANDIES achieved an ARI score 14% higher than that of SEDR, demonstrating CANDIES’ superior denoising capability through the integration of the protein modality information. Besides, CANDIES and SEDR achieved higher ARI scores compared to other ST analysis methods, indicating that denoising ST data indeed enhances spatial domain identification. A box plot summarizing the performance of all methods across all evaluation metrics over thirty independent trials is shown in Fig. 2c, suggesting that CANDIES stably outperformed compared methods across different evaluation metrics. As compared to SEDR and SpaGCN, CANDIES’ performance had little variability across different trials, indicating its high accuracy and robustness. To evaluate the impact of data quality on CANDIES denoising performance, we examined the ARI scores under increasing levels of Gaussian noise (Fig. 2d, left) and dropout rates (Fig. 2d, right) in RNA modality. The performance of all exiting methods degraded substantially as the noise level increased. However, by effectively leveraging the higher-quality protein modality, CANDIES exhibited a smaller performance decline compared to other methods, maintaining a superior and robust performance even under high-noise conditions. As a comparison, although SEDR performed well under low noise levels and dropout rates, its accuracy dropped dramatically as data quality deteriorated.

**Fig. 2:**
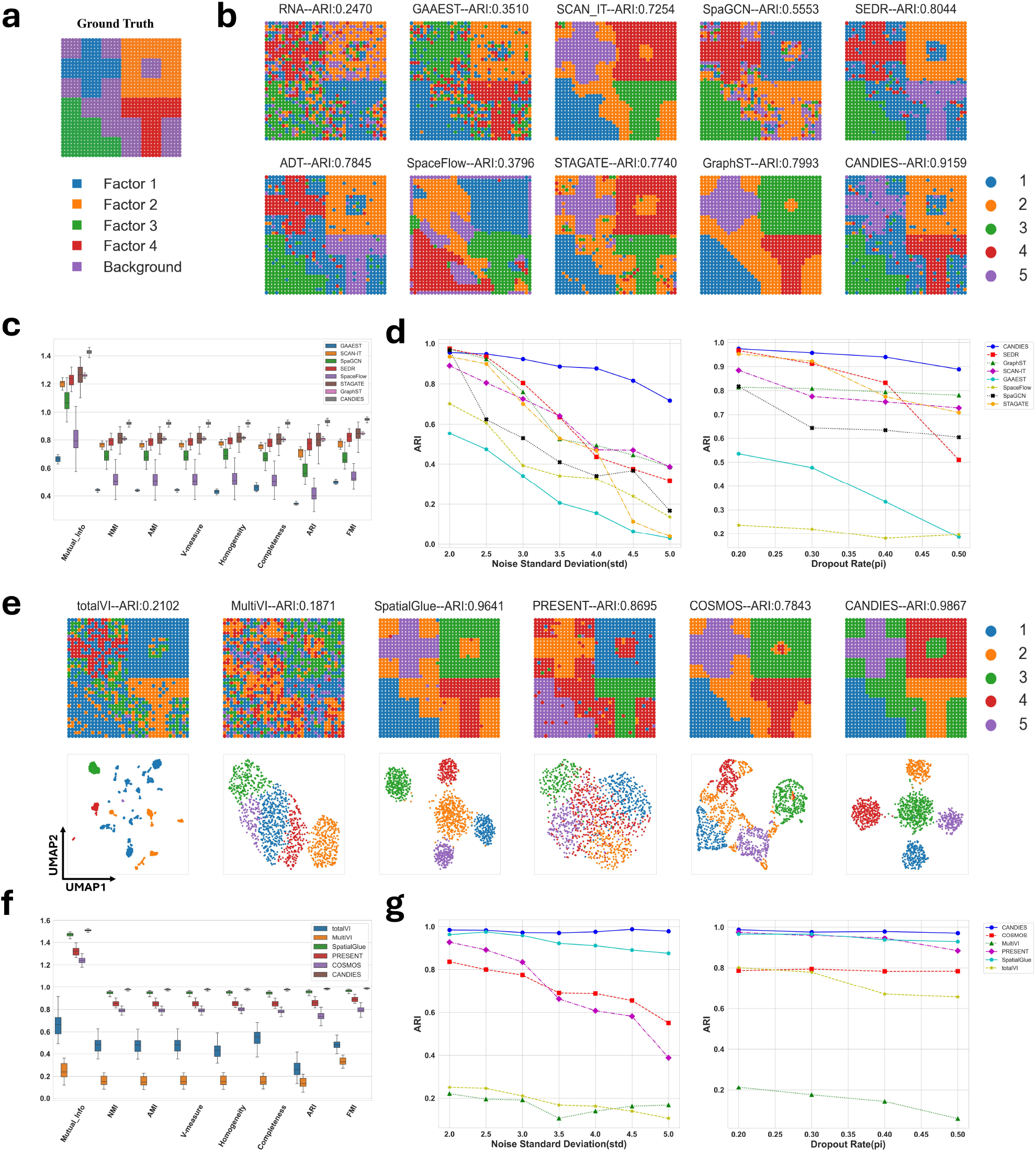
CANDIES accurately identified spatial domains in simulated spatial multi-omics dataset. **a**, Ground truth of the simulated spatial multi-omics data. **b**, Spatial plots of the simulated data with only RNA modality across eight methods designed for spatial transcriptomics (ST) data, the first column is identified by Leiden [35] on each modality. **c**, Box plots of the eight supervised metrics across the eight ST analysis methods. **d**, ARI scores with varying noise levels across eight ST methods. **e**, Spatial plots and UMAP visualization of the simulated data across six single-cell and spatial multi-omics integration methods. **f**, Box plots of the eight supervised metrics across six methods. **g**, The impact of varying noise levels on ARI across six single-cell and spatial multi-omics integration methods.

With the promising results of denoising, we assessed the performance of CANDIES in multi-omics integration by comparing it with several representative methods, including two non-spatial multi-omics methods: totalVI [15] and MultiVI [14], and three recently proposed spatial multi-omics methods, SpatialGlue [25], PRESENT [26] and COSMOS [27]. Under a moderate noise level (i.e., Gaussian noise standard deviation of 3.0 and dropout rate of 0.3), CANDIES achieved the highest ARI score of 0.9895, successfully separating all factors without any overlap (Fig. 2e). Such results demonstrated its superior integration capability. In particular, compared with the domain identification results in Fig. 2b, which was only based on the refined RNA modality, CANDIES achieved higher improvement by integrating the high-quality modality and the refined low-quality modality. As can be seen, CANDIES consistently outperformed the multi-omics methods across various metrics (Fig. 2f), demonstrating stable integration performance. Notably, spatial multi-omics methods generally achieved better results with lower variance, suggesting that incorporating the tissue microenvironment plays an important role in capturing biologically meaningful structures. Moreover, CANDIES outperformed other multi-omics integration methods that do not incorporate a denoising step, suggesting the importance of eliminating irrelevant noise while preserving biologically meaningful signals. When the noise level in the RNA modality was varied (Fig. 2g), the inconsistency in data quality across modalities significantly affected the performance of existing spatial multi-omics methods. In contrast, CANDIES consistently achieved the highest and most stable ARI score under all conditions, underscoring the critical role of denoising in spatial multi-omics analysis.

### CANDIES captures fine-grained spatial domain with denoising on spatial transcriptome-proteome data of a human skin biopsy from a COVID-19 vaccine injection site

We applied CANDIES to analyze real spatial co-profiling data of transcriptome and proteome obtained from skin biopsy tissue after COVID-19 mRNA vaccination generated by spatial CITE-seq technique [3]. The tissue section contains a collagen-rich dermis region with low cell density and a pilosebaceous unit characterized by high cellularity (Fig. 3a). The competition between ADTs and mRNAs for in-tissue reverse transcription reduces the efficiency of transcript detection [3], making the RNA modality highly noisy. Therefore, although the overall structures of the dermis and pilosebaceous units were clearly discernible in the protein modality, these anatomical features appeared less distinguishable in the RNA modality (Fig. 3b). The inherent noise in the RNA modality made it difficult to reliably identify biological clusters with existing methods (Fig. 3b). For example, SCAN-IT, SpaGCN, GraphST, and STAGATE exhibited significant noise and failed to clearly identify the major regions. In contrast, CANDIES effectively leveraged the high-quality protein modality for denoising, enabling accurate identification of the dermis and the pilosebaceous unit. This result demonstrated the superiority of CANDIES in handling the highly noisy ST data.

**Fig. 3:**
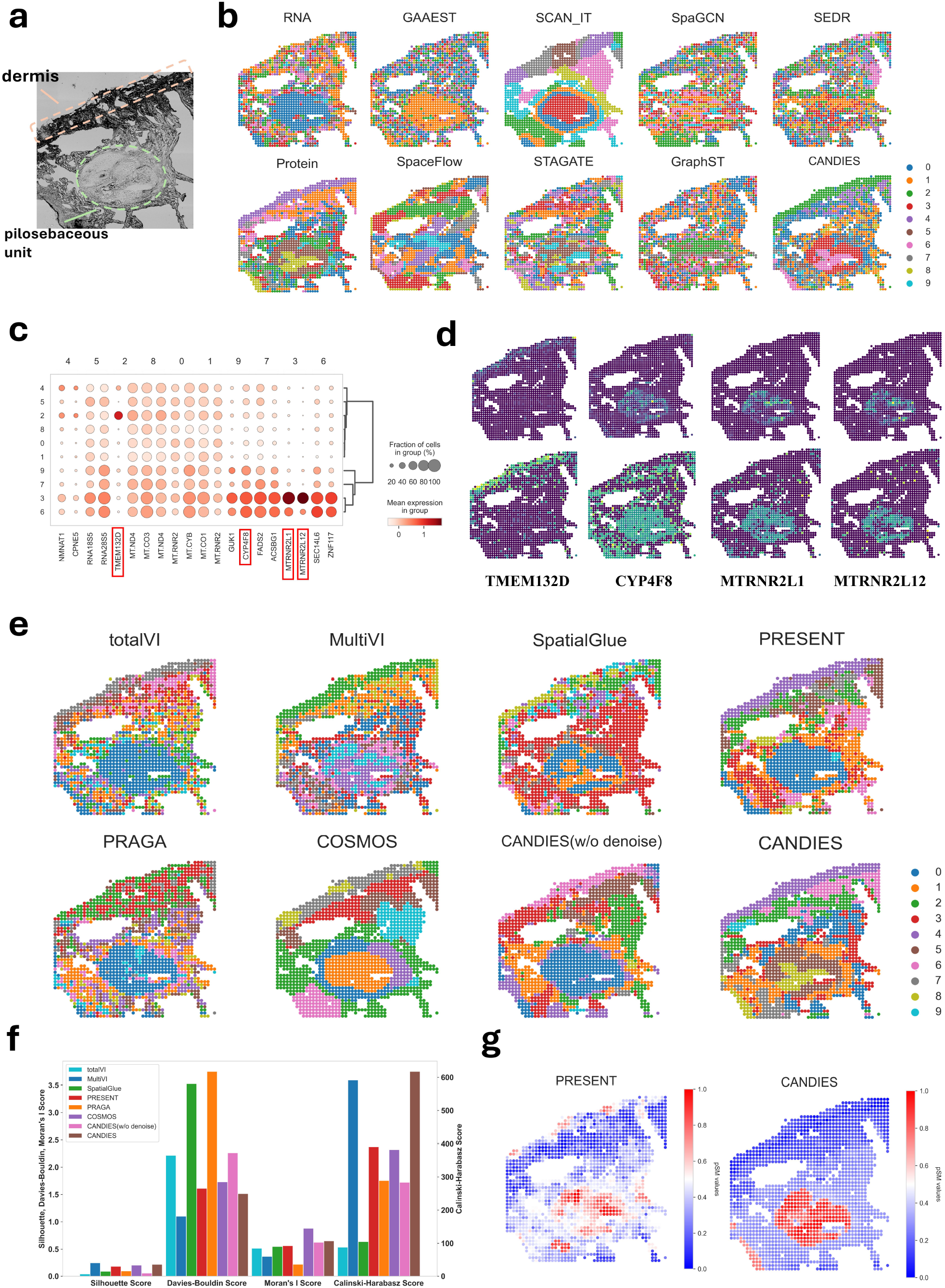
CANDIES achieves fine-grained spatial partitioning on the human skin dataset generated using the spatial CITE-seq technique. **a**, The bright-field image of the human skin tissue section. Highlighted regions are dermis and pilosebaceous unit. **b**, Spatial plots of the human skin data only with RNA modality across eight representative spatial transcriptomics (ST) methods, the first column is identified by Leiden on individual modality. **c**, Top-2 differentially expressed genes (DEGs) identified for each cluster discovered by CANDIES on the denoised ST dataset. The dot size represents the proportion of spots within a cluster that express the corresponding DEG, and the color intensity denotes the average expression level of that DEG in the cluster. **d**, The first and second rows correspond to spatial distribution of the selected four DEGs before and after denoising. **e**, Spatial plots of the human skin data across seven single-cell and spatial multi-omics integration methods, CANDIES (w/o denoise) refers to the variant of CANDIES without the denoising phase. **f**, Bar plots comparing the performance of seven methods across four unsupervised metrics. Note that a lower Davies-Bouldin Index (DBI) indicates better clustering performance, whereas higher values on the other metrics reflect improved results. **g**, Pseudo spatiotemporal maps (pSM) generated by CANDIES and PRESENT.

We further examined the spatial expression patterns of differentially expressed genes (DEGs) before and after denoising (Fig. 3c-d). Among the identified top-2 DEGs, we found multiple lines of evidence aligned with prior knowledge. Consistent with original study [3], the transmembrane protein-encoding gene *TMEM132D* in cluster 2 showed specific expression in the dermis region, while the *CYP450* protein-encoding gene *CYP4F8* were broadly expressed across the most skin regions. Additionally, *MTRNR2L1* and *MTRNR2L12* showed specific enrichment in the pilosebaceous unit (cluster 3 and cluster 6). Notably, the spatial expression patterns of the four selected DEGs became substantially more distinct in the denoised RNA modality compared to their original profiles, demonstrating that CANDIES’ denoise step effectively captures meaningful biological signals.

Next, we comprehensively benchmarked CANDIES’ integration performance with totalVI [15], MultiVI [14], SpatialGlue [25], PRESENT [26], PRAGA [28] and COSMOS [27] on spatial transcriptome-proteome human skin data (Fig. 3e). We also included a simplified version of CANDIES that takes the original modalities as input, allowing us to investigate the impact of denoising step. Without incorporating the spatial information, TotalVI and MultiVI produced less smooth boundaries between spatial domains. Meanwhile, spatial multi-omics methods such as SpatialGlue and PRAGA failed to depict the dermal architecture, since they do not consider the huge differences of data quality across modalities. PRESENT showed relatively better performance, successfully identifying the main regions, e.g., the dermis region and the pilosebaceous unit, but still exhibited limitations in handling fine-grained details. The clustering patterns produced by CANDIES more closely resembled those identified on individual modality. For example, cluster 8 was only identified by CANDIES but remained undetected by PRESENT, MultiVI, and the simplified version of CANDIES without denoising, validating the effectiveness of denoising in distinguishing fine-grained biological patterns. We employed unsupervised metrics to quantitatively evaluate the clustering performance. CANDIES outperformed all other methods with the highest Calinski-Harabasz Score and ranked among the top two methods in terms of Silhouette Score, Moran’s I, and Davies-Bouldin Score, highlighting the importance of denoising in enhancing spatial multi-omics integration.

To investigate the spatiotemporal dynamics across different regions, we leveraged CANDIES clustering results to generate a pseudo-Spatiotemporal Map (pSM) for the skin biopsy [36] (Fig. 3g), using PRESENT as a baseline for comparison. We observed that the pSM values of CANDIES were lowest in the dermal region and exhibited a gradual increase toward the pilosebaceous unit, reflecting a plausible spatial developmental trajectory from dermal to pilosebaceous. This spatial gradient in pSM values aligns with the known developmental sequence of these structures, where the dermis serves as the origin for sweat glands, sebaceous glands, apocrine glands, and hair follicles [37]. In contrast, PRESENT produced pSM values with a very noisy dermal-to-pilosebaceous gradient, likely due to the intrinsic noise in RNA modality.

### CANDIES facilitates the integrated analysis of spatial epigenome-transcriptome data from mouse brain and mouse embryo

Next, we applied CANDIES to integrate the spatially resolved epigenome-transcriptome datasets to showcase its ability to handle different types of omics data. To demonstrate the broad applicability of CANDIES to a wide spectrum of sequencing platforms, we considered two spatially resolved epigenome-transcriptome datasets, mouse brain data generated with MISAR-seq [7] and mouse embryo data generated with spatial ATAC–RNA–seq [6], representing different noise patterns introduced by different sequencing technologies.

We first applied CANDIES to a mouse brain dataset generated using MISAR-seq platform [7], which includes eight sequencing datasets from four key developmental stages: E11.0, E13.5, E15.5, and E18.5. In this section, we focused on the E18.5 mouse brain data. First, we compared CANDIES with the spatial multi-omics integration methods SpatialGlue, PRESENT, and COSMOS, as well as the single-cell methods MultiVI and scMDC [38] (the denoising results of CANDIES were shown in Supplementary Fig. S3). TotalVI was not included as it is specifically designed for single-cell RNA-ADT data. To determine the relative quality of the two modalities, we applied Leiden method on the low-dimensional embeddings of each modality obtained from a pre-trained graph auto-encoder and evaluated the ARI scores based on the manual annotation collected from Kaufman’s Atlas of Mouse Development and Allen Brain Atlas [7] (Fig. 4a). As shown in the first column of Fig. 4b, it was clear that the ATAC modality had better performance in capturing regions such as the mantle zone of the dorsal pallium (DPallm), cartilage_1, and cartilage_2, achieving a higher ARI score. Therefore, we used the ATAC modality as the condition to refine the quality of the RNA modality in the denoising step of CANDIES.

**Fig. 4:**
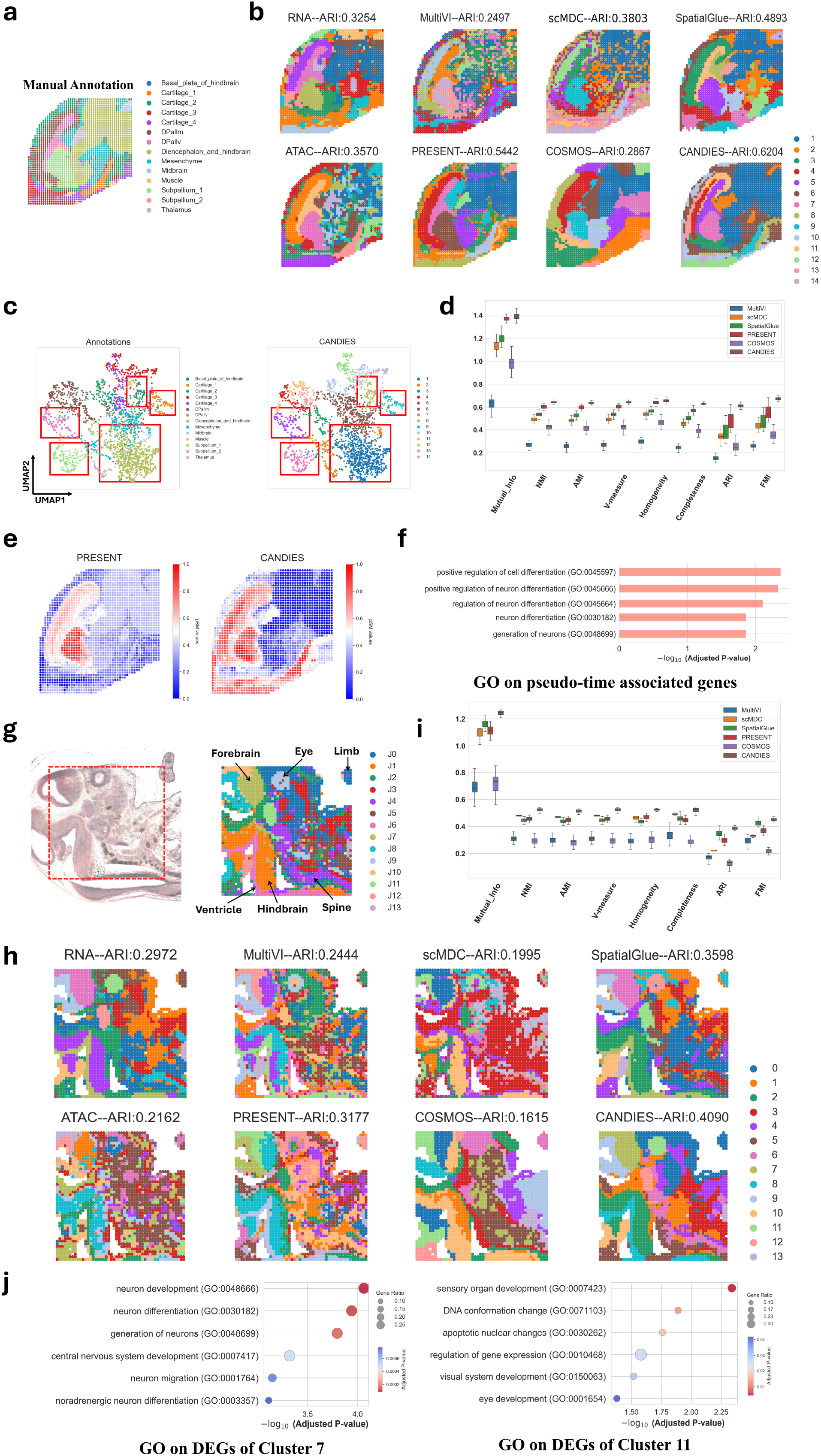
CANDIES accurately integrated spatial multi-omics data from the mouse brain and mouse embryo. **a**, Manual annotation of the E18.5 mouse brain data. **b**, Spatial plots of the E18.5 mouse brain data across six single-cell and spatial multi-omics integration methods, the first column is identified by Leiden on each modality. **c**, The UMAP visualization of CANDIES’ latent representations colored by manual annotation (left) and CANDIES (right). **d**, Box plots of the eight supervised metrics across the six methods. **e**, Pseudo spatiotemporal maps (pSM) generated by CANDIES and PRESENT. **f**, Gene Ontology Biological Process (GOBP) enrichment analysis of pseudo-time associated genes. **g**, The bright-field image (left) and the manual annotation (right) of the E13 mouse embryo data. **h**, Spatial plots of the E13 mouse embryo data across six single-cell and spatial multi-omics integration methods, the first column is identified by Leiden on each individual modality. **i**, Box plots of the eight supervised metrics across the six methods. **j**, Gene Ontology (GO) enrichment analysis on differentially expressed genes (DEGs) of cluster 7 (left) and cluster 11 (right).

As shown in Fig. 4b, we observed that CANDIES demonstrated superior results in both major domain identification and the preservation of rare spot types. Specifically, CANDIES successfully distinguished major anatomical categories, including cerebral regions (diencephalon_and_hindbrain, cluster 1), mesenchyme (cluster 6), and the subpallium subtype (subpallium_1, cluster 7), while also preserving the integrity of rare types such as the ventricular zone of the dorsal pallium (DPallv, cluster 5) and two cartilage subtypes (cartilage_1 and cartilage_2, clusters 8 and 9). In contrast, other methods exhibited notable limitations. For example, since single-cell multi-omics methods such as MultiVI and scMDC do not incorporate spatial information, their clustering results exhibited poor spatial continuity. This led to fragmented spatial distributions, where adjacent regions were often incorrectly assigned to different clusters, failing to accurately represent the true tissue organization. Consistently, these methods also received the lowest scores in supervised evaluation metrics, further supporting this conclusion (Fig. 4d). Spatial multi-omics methods demonstrated improved performance; however, their inability to effectively handle noise sometimes led to incorrect partitioning of regions. For instance, PRESENT successfully identified DPallm, DPallv, and subpallium_1. But it erroneously divided the Diencephalon_and_hindbrain into two separate regions. Similarly, SpatialGlue showed moderate performance, correctly identifying subpallium_1, DPallm, and DPallv, yet wrongly clustered the spots within the diencephalon_and_hindbrain region, fragmenting them into multiple smaller regions that did not align with known anatomical structures. In contrast, CANDIES consistently achieved higher scores with lower variance (Fig. 4d), indicating its superior capability to integrate epigenome-transcriptome data through cross-modality denoising. As complementary evidence, the UMAP visualization of spot embeddings generated by CANDIES further verified its ability to effectively distinguish key regions, including the cerebral regions (diencephalon_and_hindbrain), mesenchyme, and cartilage_2 (Fig. 4c). This distinction is particularly important given the functional and developmental importance of these regions. For instance, the diencephalon_and_hindbrain play distinct roles in neural development and function, making their accurate identification crucial for understanding brain organization [39]. Similarly, differentiating mesenchyme from cartilage subtypes is essential for studying skeletal development and associated pathologies [40]. These findings underscore the robustness of CANDIES in preserving biologically meaningful spatial structures within complex spatial epigenome-transcriptome data.

To investigate the spatiotemporal patterns of the brain omics data, we derived pSM values using the clustering results from PRESENT and CANDIES (Fig. 4e). Both CANDIES and PRESENT revealed clear and smooth layered structures. However, the pSM values derived from CANDIES not only aligned with the overall spatial organization of the E18.5 mouse brain, but also showed strong concordance with manually annotated anatomical boundaries, particularly in several critical regions such as the diencephalon, hindbrain, subpallium, and cartilage. To further validate these findings, we identified 102 pseudo-time-associated genes based on the pSM values calculated with CANDIES embeddings. Gene Ontology (GO) enrichment analysis revealed that these genes were significantly enriched in biological processes related to cell differentiation, neuron differentiation and generation of neurons. These results underscore the ability of CANDIES to maintain biological variations, providing a robust framework for studying complex developmental processes in the brain.

Furthermore, we applied CANDIES to integrate epigenome-transcriptome data of generated with E13 mouse embryo data derived from spatial ATAC–RNA–seq [6]. Based on the spatial ATAC–RNA–seq description, we manually annotated the major regions in this data (Fig. 4g, right), including the forebrain, eye, limb, ventricle, hindbrain, and spine. Unlike the MISAR-seq dataset, we observed that the ARI score of the RNA modality was greater than that of ATAC modality, suggesting a better quality of the transcriptome profiles (Fig. 4h, first column), which was consistent with the original study [6]. Therefore, we utilized the RNA modality to enhance the ATAC modality in this dataset. As shown in Fig. 4h, CANDIES produced the most accurate spatial domain identification among compared methods. MultiVI, SpatialGlue, and PRESENT generated less distinct clustering patterns with substantial overlap across different domains. In contrast, CANDIES effectively identified all major anatomical regions including hindbrain (cluster 2), ventricle (cluster 8), and eye tissues (cluster 11), achieving the highest ARI score of 0.4090, which is 13.7% higher than the runner-up. Importantly, the eye region was not detected by other spatial multi-omics methods, likely due to the noisy ATAC modality. CANDIES exclusively identified this region by taking the advantage of the denoising step. Other supervised metrics also indicated that CANDIES consistently outperformed existing methods in terms of domain identification (Fig. 4i).

Finally, to characterize the biological significance of the domains identified by CANDIES, we performed GO enrichment analysis (Fig. 4j) with differentially expressed in two regions: cluster 7 (annotated as forebrain) and cluster 11 (annotated as eye). For each cluster, we identified 20 differentially expressed genes (DEGs), and performed GO enrichment analysis. We identified significant enrichment that aligns with the biological function of the corresponding clusters. Specifically, cluster 7, corresponding to forebrain, showed pronounced enrichment in fundamental neuro-developmental processes, including generation of neurons, neuron differentiation, neuron development and neuron migration, which are closely related to the forebrain’s well-established role in neurogenesis and cortical circuit formation [41]. For cluster 11, we found it strongly associated with the eye development process and exhibited enrichment for sensory system development pathways, particularly sensor organ development and visual system development. These cluster-specific GO enrichment results suggest that CANDIES successfully extracted biologically meaningful signals by integrating ATAC–RNA–seq data.

### CANDIES improves the mapping of complex traits to relevant spatial domains

Although the advancement of spatial omics has provided valuable insight into the spatial landscape of a cascade of biological process across a wide range of tissues, most of the available spatial data resources are not pertinent to a specific trait of interest. Integrating spatial omics data with complex trait GWASs is expected to offer a comprehensive characterization of how the spatial distribution of cells are associated to disease pathology. However, the strong noise present in spatial omics data makes it difficult to establish a reliable trait-spot link. In this section, we show that our spatially informed cross-modality embeddings produced by CANDIES can improve the mapping of trait-spot associations. To illustrate this benefit, we carried out gsMap [42] analysis by taking the embeddings of CANDIES as input to identify the trait-spot associations across 3 tissues, including mouse brain, mouse embryo, and human lymph, and 32 complex traits encompassing 5 broad biological clusters. Briefly, to link polygenic GWAS signals to spatial locations, we define each spot as an annotation of a set of single nucleotide polymorphisms (SNPs) based on its gene specificity score (GSS), and estimate the heritability enrichment of each spot for a target trait by employing the stratified linkage disequilibrium score regression (S-LDSC). Since each spot represents a noisy observation of the true spatial profile, we shall aggregate the information across homogeneous spots to obtain a reliable annotation. Here, we first applied CANDIES to combine multi-omics spatial data, leading to a more robust latent representation of the original profile. Then, a homogeneous microdomain was constructed for each spot based on the cosine similarity of the CANDIES latent embeddings. Such a microdomain includes a set of spots sharing spatial and biological similarities with the focal spot across different modalities. For each gene, we estimated its GSS within each focal spot by calculating a normalized geometric mean of its expression rank across the microdomain of the focal spot. Finally, we performed S-LDSC analysis by using each spot’s GSS as the annotation. As such, the trait-associated spots can be identified by testing whether SNPs with higher GSS are enriched for heritability.

We summarized the identified trait-associated spatial locations in Fig. 5 and Supplementary Figs. S7-S14. Fig. 5 shows the *p*-values of enrichment across different spatial regions, obtained by aggregating spot-level *p*-values using Cauchy combination test. As we can observe, the enriched regions were highly aligned with our knowledge of tissue regions related to the 32 traits. For psychiatric traits, such as bipolar disorder (BIP), intelligence (IQ), major depression disorder (MDD), schizophrenia (SCZ), and behavioral phenotypes, including education attainment (EA) and smoking initiation (SmokingI), we observed significant enrichment for heritability in the dorsal pallium, thalamus, and hindbrain of the mouse brain data (Figs. 5a-c, Supplementary Figs. S7a-b and Supplementary Fig. S8). In mouse embryo data (Fig. 5d), psychiatric and behavioral traits were also strongly associated with regions related to central nervous system (CNS), which is consistent with our findings in the mouse brain data (Fig. 5e and Supplementary Fig. S7c). Besides, height was mapped to embryo regions related to the formation of cartilage (Fig. 5f). In the analysis of lymph node data, we found substantial enrichment of haemotological traits within regions of cortex, follicle, and capsule (Fig. 5g-i, Supplementary Fig. S7e-f and Supplementary Fig. S12). The three lines of evidence from different spatial multi-omics data converged to well-established biological relevance between traits and tissues, suggesting CANDIES embeddings successfully captured important biological information to facilitate interpreting GWAS discoveries. Due to the noisy nature of RNA modality, the trait-relevant regions were less enriched when the embeddings were generated with only RNA modality (Supplementary Fig. S7e). Given the promising results, we investigated the spatial distribution of trait associations at the spot level. We found that cells located closer to the dorsal end of the dorsal pallium exhibited a much stronger association with MDD (Fig. 5b) and SCZ (Fig. 5c). Previous studies have reported that such observation can be attributed to the spatially differential expression of calcium transport-related genes in Glu-neurons along the dorsal-ventral axis [42]. Therefore, by integrating with GWAS data, the joint embeddings produced by CANDIES can offer a better understanding of the genetic architecture of complex human traits within spatially resolved tissue profiles.

**Fig. 5:**
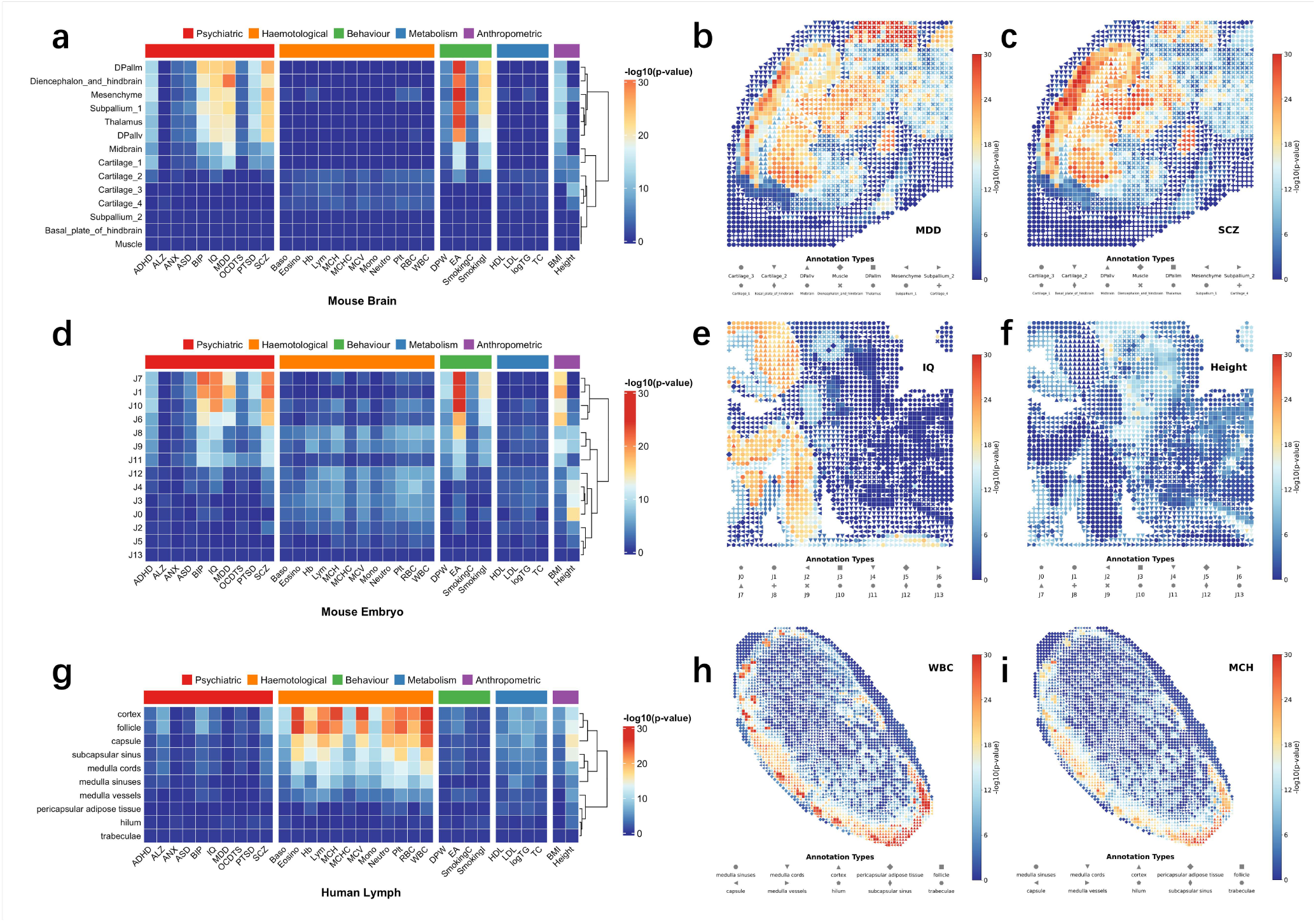
Trait-spot associations identified by using CANDIES embeddings in three spatial omics datasets as input. **a**,**d**,**g**, Heatmaps showing the significance level of associations between complex traits and spatial regions, with rows representing annotated regions in the mouse brain, mouse embryo, and human lymph respectively, and columns representing human traits categorized into 5 classes. *p*-values are obtained with Cauchy combination test across spots within each region. Color scale indicates the − log_10_ *p*-values of the Cauchy combination test. **b**,**c**, Heritability enrichment of two representative psychiatric traits, major depressive disorder and schizophrenia, in the mouse brain analysis, with spot annotation indicated by different point shapes and − log_10_ *p*-values reflected by the color. **e**,**f**, Heritability enrichment of two representative traits, IQ and height, in the mouse embryo analysis, illustrating how developmental regions associate with cognition and anthropometric traits. Annotations J0-J13 is obtained from the joint clustering of spatial ATAC and RNA data, where J1 and J6 roughly map to the Hindbrain region, J7 and J10 map to the Forebrain region, J3 and J4 map to the spine, and the circular area marked by J9 maps to the eye region. **h**,**i**, Heritability enrichment of two hematological traits, white blood cells count and mean corpuscular hemoglobin, in the human lymph analysis.

## Discussion

In this paper, we have introduced CANDIES, a novel multi-omics integration framework based on conditional diffusion and contrastive learning-based framework. CANDIES is able to integrate two modalities with varying qualities, thereby generating robust and unified spatial representations. Through extensive experiments, we showed that CANDIES outperforms top-performing methods, including 7 spatial single-omic clustering methods, 3 single-cell multi-omics integration approaches, and 4 spatial multi-omics integration techniques. We applied CANDIES to integrate spatially resolved multi-omics data using four real datasets comprising varying tissue types, sequencing platforms, and noise patterns, producing high-quality low-dimensional representations that are versatile in multiple down-stream biological analysis. We showed that CANDIES embeddings are highly reliable in denoising low-quality modality, spatial domain identification and pseudo-spatiotemporal map generation. By combing the output of CANDIES with complex trait GWASs, we illustrated that the output of CANDIES was particularly helpful to characterize the spatially-resolved mapping of complex human traits. We believe that CANDIES will serve as a powerful analytic tool to facilitate spatial multi-omics data analysis.

The key strength of CANDIES lies in its two-stage framework, which first refines the lower-quality modality using a conditional diffusion model and then integrates the enhanced data with another modality through contrastive learning. Unlike conventional spatial multi-omics integration methods, CANDIES leverages guidance from the higher-quality modality and spatial information based on DiT network to achieve effective noise reduction for the lower-quality modality. During integration, CANDIES first employs graph auto-encoders to derive both spatial and feature embeddings for each modality. Then, an innovative contrastive learning strategy is applied to align feature or spatial embeddings across different modalities. Finally, an attention mechanism dynamically balances spatial and feature information. This systematic approach not only enhances robustness and preserves cellular heterogeneity but also guarantees biologically meaningful results, positioning CANDIES as a uniquely effective solution for diverse spatial multi-omics datasets.

In the future, there are several promising avenues for extending CANDIES. One potential direction is the incorporation of imaging data as an additional modality [43, 44]. Many spatial multi-omics technologies produce accompanying imaging data, such as H&E-stained images, which offer critical information about cell and tissue morphology. We plan to extend CANDIES to integrate image data, either during the denoising phase or the integration phase, further enhancing its analytical capabilities. Additionally, we aim to expand CANDIES to support triple-omics and single-cell level analysis, addressing the growing complexity of multi-omics datasets and enabling more comprehensive biological insights.

## Methods

### Data description and preprocessing

To comprehensively evaluate the performance of CANDIES, we conducted extensive quantitative and qualitative experiments across a simulated spatial multi-omics dataset, and four real-word spatially resolved multi-omics datasets.

#### Simulated Data

To evaluate the spatial multi-omics data analysis methods, we first followed SpatialGlue [25] to generate paired spatial multi-omics datasets comprising two modalities (RNA and protein). Within it, the ‘ggblocks’ model [45] was employed to produce the spatial pattern with four different factors of 1296 spots (or cells). For the RNA modality, the gene expression matrix **X**^*rna*^ ∈ ℝ^1296×800^ was generated from a Zero Inflated Negative Binomial (ZINB) distribution with the mean defined as: **M**^*rna*^ = **B**^*rna*^ + **F**^*rna*^**W**^*rna*^, where the value of every element in **B**^*rna*^ was 0.5, representing the mean value of background expression. **F**^*rna*^ ∈ ℝ^1296×4^ was a binary matrix indicating the activity of spatial factors at each spot, 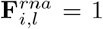 if the spatial factor *l* was active at spatial spot *i*, otherwise 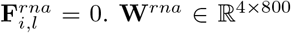 was the mean expression matrix for spatial factors. The dropout rate in ZINB distribution was set to pi, and the shape parameter for the negative binomial distribution was set to 3. For the protein modality, the protein expression matrix **X**^*pro*^ ∈ ℝ^1296×100^ drawn from a negative binomial distribution with the mean defined as: **M**^*pro*^ = **B**^*pro*^ + **F**^*pro*^**W**^*pro*^, where each element in **B**^*pro*^ was 2, **F**^*pro*^ ∈ ℝ^1296×4^ and **W**^*pro*^ ∈ ℝ^4×100^ were configured analogously to the RNA modality. The shape parameter was also set to 3.

To further simulate the quality differences between modalities, we introduced a higher level of Gaussian noise to the RNA modality and inflated its drop-out rate. Specifically, for the RNA modality, Gaussian noise with a mean of 2 and standard deviations ranging from 3 to 6, increasing in steps of 0.5, was introduced to the expression matrix **X**^*rna*^. Furthermore, we also varied the dropout rate from 0.2 to 0.5 with an increment of 0.1. For the protein modality, we introduced a lower level of Gaussian noise with a mean of 2 and a fixed standard deviation of 1 on **X**^*pro*^. The detailed parameters are shown in the Supplementary Tables. S3-S4.

#### Spatial CITE-seq human skin dataset

Spatial CITE-seq was employed to map early immune cell activation within a skin biopsy tissue collected from the injection site of a Coronavirus Disease 2019 (COVID-19) mRNA vaccine, co-profiling spatially resolved proteome and transcriptome [3]. Unlike previous study, which analyzed a much smaller set of proteins and struggled with tissue region clustering based solely on protein profiles, spatial CITE-seq employs a cocktail of approximately 200–300 antibody-derived tags (ADTs) to stain a tissue slide. This dataset comprises 15,486 genes and 283 proteins on 1691 spots.

For the RNA modality, we first removed genes expressed in fewer than 10 spots to eliminate low-quality features. Subsequently, we normalized the gene expression count using the *scanpy*.*pp*.*normalize total* function, followed by a log transformation and standardization to ensure comparability across cells. Finally, we identified the top 3000 highly variable genes (HVGs) using the Seurat v3 [46] implemented in scanpy package (v.1.10.3) [47]. For the protein modality, we employed the centered log-ratio (CLR) transformation to normalize the count data for each spot [48]. Following CLR transformation, we standardized the data using the *scanpy*.*pp*.*scale* function.

#### MISAR-seq E18.5 mouse brain dataset

MISAR-seq enables the simultaneous profiling of chromatin accessibility and gene expression during mouse brain development [7]. E18.5 mouse brain dataset comprises 2,129 spots, 32,285 genes, and a peak count ranging up to 161,461.

For the RNA modality, we removed genes expressed in fewer than 20 spots and selected spots that contain more than 100 expressed genes. Then, we normalized the gene expression count followed by a log transformation. Finally, we identified the top 4,000 HVGs by scanpy package (v.1.10.3) [47]. For the ATAC modality, as the dimensions are much higher than that of the RNA modality, we applied Latent Semantic Indexing (LSI) to select informative features [49, 50]. Specifically, we first used TF-IDF (Term Frequency-Inverse Document Frequency) normalization, and the normalized data were log-transformed and further processed with randomized Singular Value Decomposition (SVD) to reduce dimensionality and capture the most meaningful features in the count data.

#### Spatial ATAC–RNA-seq E13 mouse embryo dataset

Spatial ATAC-RNA-seq enables spatially resolved, genome-wide co-mapping of the epigenome and transcriptome by simultaneously profiling chromatin accessibility and messenger RNA expression [6]. In experiments conducted on embryonic day 13 (E13) mouse embryos, with a pixel size of 50 *µm*, the dataset comprises 2,186 spots, 20,900 genes, and a peak count reaching up to 87,173, offering a detailed multi-omics view of embryonic development.

For the RNA modality, we removed genes expressed in fewer than 20 spots, then normalized and log-transformed the gene expression count. Finally, we identified the top 3,000 HVGs using the scanpy package (v.1.10.3) [47]. For the ATAC modality, we performed preprocessing steps similar as the E18.5 mouse brain dataset.

#### 10x Visium human lymph node dataset

The human lymph node dataset was generated using 10x Genomics Visium spatial RNA-protein co-profiling technology. This spatially resolved multi-omics dataset originates from a comprehensive study focused on multi-modal representation learning, capturing both transcriptomic and proteomic profiles within their native tissue architecture. The dataset contains 3,484 spots, 18,085 genes, and 31 proteins.

For the RNA modality, we first filtered out low-quality features by removing genes expressed in fewer than 20 spots. The remaining gene expression counts were then normalized and followed by a log transformation. To enable cross-sample comparisons, we standardized the transformed data. Finally, we identified the top 3000 highly variable genes (HVGs). For the protein modality, we performed the same preprocessing workflow as the human skin dataset.

### The CANDIES model

CANDIES utilizes the powerful generative capabilities of diffusion models to refine the modality with lower quality, as described by Fig. 1, the framework of it consists of two main parts: denoising and integration. For better elaboration, we assume there are two modalities, where the quality of modality 2 is superior than that of modality 1. Denoting the expression matrix of modality 1 and modality 2 are 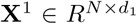 and 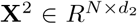 respectively, where *N* represents the number of spots (or cells) in the dataset, and *d*_*m*_ corresponds to the features dimension in the *m*-th modality, *m* ∈ {1, 2}. The spatial coordinates of these *N* sequencing spots (cells) are defined as 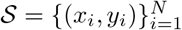.

#### Denoising phase of CANDIES

In the denoising part, a pre-trained dual graph auto-encoder is firstly employed to obtain the low-dimensional embeddings of all spots for two modalities. These embeddings are then fed into decoders to reconstruct the original expression matrix. For the embeddings of lower-quality modality, it performs as the starting point of the diffusion process, and a DiT-based conditional denoising model with the embeddings of the higher-quality modality as condition is utilized to learn the reverse process, beginning from these noise-perturbed initial embeddings.

##### Construction of spatial neighbor graph

For the given modality 1 or modality 2, we firstly utilized their spatial information to construct the spatial graph as 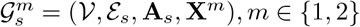, where 𝒱 denotes the set of *N* spots and ℰ_*s*_ represents the set of edges connecting these spots. **A**_*s*_ ∈ ℝ^*N* ×*N*^ is the adjacency matrix calculated by the spatial coordinates 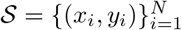, where **A**_*s*_(*i, j*) = 1 if the spots *j* is among the *k*-nearest neighbors of the spots *i* (*k* = 3 by default), otherwise **A**_*s*_(*i, j*) = 0.

##### Pre-trained graph auto-encoders (GAEs) for individual modality

To enable efficient training of diffusion model, high-dimensional features from the original data space are firstly mapped to a compact embedding space using a graph auto-encoder. Two different encoders 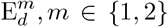, consisting of two graph convolution network (GCN) layers, are employed to generate the low-dimensional embedding 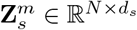 from spatial graph 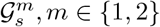 respectively. Moreover, two two-layers GCN decoders 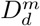 are employed to reconstruct the features for two modalities from these latent embeddings. Mathematically, the reconstruction process is expressed as follows:

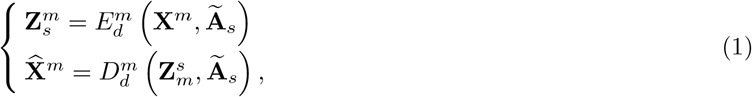

where **X**^*m*^ and 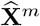 represent the original expression matrix and reconstructed expression matrix of modality *m*. 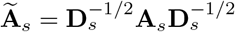 is the normalized adjacency matrix, **D**_*s*_ is the degree matrix of **A**_*s*_.

The reconstruction loss ℒ_rec1_ is computed to measure the discrepancy between the original input features **X**^*m*^ and the reconstructed features 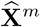:

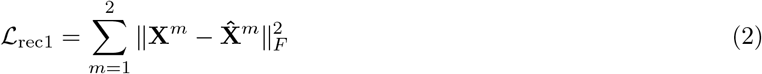

It should be noted that a Zero Inflated Negative Binomial (ZINB) [51] decoder is employed for the RNA modality to address the prevalent dropout events, which arise from both technical and biological factors. Specifically, the ZINB decoder contains three separate fully connected layers positioned after two-layers GCN, which is designed to estimate the three key parameters of the ZINB distribution, including the dropout rate ***π***, the dispersion parameter ***θ***, and the mean ***µ***:

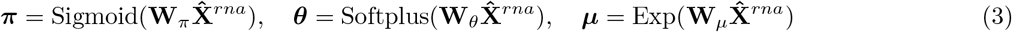

where 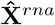 represents the reconstructed gene expression matrix generated by Eq. (1), **W**_*π*_, **W**_*θ*_ and **W**_*µ*_ are learnable weight matrices of fully connected layers. Sigmoid(), Softplus() and Exp() denote the sigmoid, soft-plus and exponential function separately. Then the expression of the *j*-th gene on the *i*-th spot satisfies the ZINB distribution: 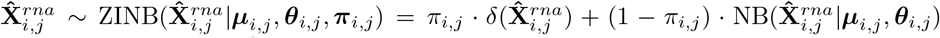 where *δ*(*x*) is the Dirac delta function, and 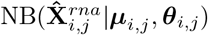 is the Negative Binomial (NB) distribution defined as: 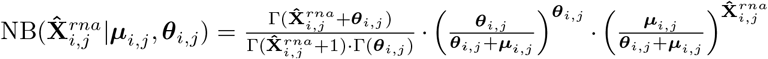, where Γ() is the Gamma function. Therefore, when one of the modality is RNA, to capture the sparse and overdispersed distribution of RNA accurately, the following reconstruction loss is employed as:

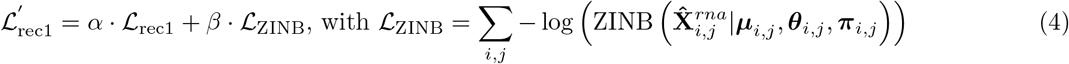

where the coefficients *α* = 2 and *β* = 0.1 are adopted in the experiment.

By optimizing Eq. (2) in the absence of RNA or Eq. (4) when RNA is one of the modalities, we derive the low-dimensional embeddings 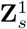 and 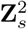 for the two respective modalities.

##### Training process of conditional diffusion model

A conditional diffusion model [52, 53] is designed to leverage the embeddings of the higher-quality modality to guide the refinement of the lower-quality modality, ensuring a more accurate and biologically meaningful generation. For concise illustration, we denote the low-dimensional embeddings of the lower-quality and higher-quality modalities as 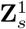 and 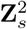, respectively. The diffusion model involves two main processes: the forward and reverse processes. In the forward process, 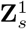 is initialized as the starting point **Z**_0_, and it is gradually transferred to pure Gaussian noise **Z**_*T*_ based on the conditional probability 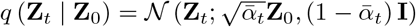. By applying the reparameterization trick, we can derive the noisy features **Z**_*t*_ at the *t*-th step:

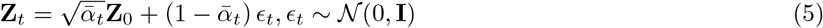

Here, 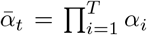, and a cosine noise scheduling mechanism is employed to enable smooth noise distribution, that is:

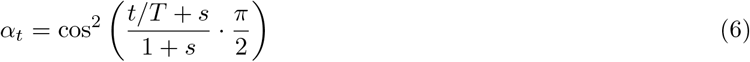

where the hyper-parameter *s* ∈ (0, 1) is used to control the noise level at each iteration.

As shown in Fig. 1, the reverse process begins from the input embeddings constructed by concatenating the two modalities 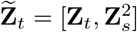. Besides, the embeddings of the higher-quality modality 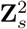 are processed by a Multi-Layer Perceptron (MLP) to obtain the representations 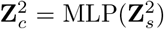, which are employed as condition to denoise the embeddings of the lower-quality modality through a learnable DiT based conditional denoising model. Diffusion Transformer (DiT) [29] utilizes transformers as the backbone of diffusion models, achieving superior performance over traditional U-Net-based approaches. When applied to spatial multi-omics data, DiT’s multi-head self-attention mechanism effectively captures both local and global spatial dependencies while also enhancing the understanding of cross-modality relationships. Specifically, the noisy features 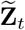 are progressively restored by the following:

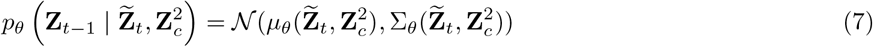

where 𝒩 represents a Gaussian distribution parameterized by the mean 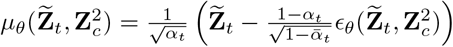 and variance 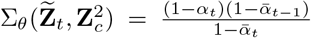, the neural network is employed to predict the statistics of *p*_*θ*_. By reparameterizing *µ*_*θ*_ as a noise prediction network *ϵ*_*θ*_, the denoising model can be optimized using the mean squared error between the predicted noise 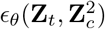 and the ground truth Gaussian noise *ϵ*_*t*_:

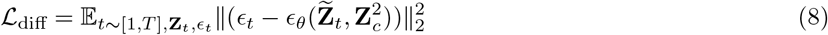

#### Inference process of conditional diffusion model

Once the DiT-based conditional diffusion model is trained, the denoised embeddings 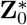 of the lower-quality modality could be enhanced by the higher-quality modality through the learned cross-modality relationships. Concretely, a random initial value 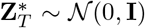 is sampled and the inference process at each time step *t* is performed by using trained DiT based conditional denoising model (Eq. (7)), i.e., 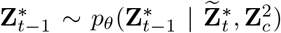, where 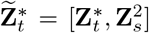. Finally, the embeddings 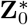 of the lower-quality modality could be iteratively recovered after *T* steps, and its corresponding expression matrix could be reconstructed through the pre-trained decoder according to Eq. (1), as formulated by 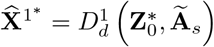.

#### Integration phase of CANDIES

After the denoising phase, the quality of spatial multi-omics data is significantly enhanced. In the integration phase, a unified representation is obtained by leveraging both the spatial information and expression data from multiple modalities. Specifically, graph convolutional networks (GCNs) are employed on spatial graph or feature graph to extract spatial or feature embeddings for every spot within each modality. Then, contrastive learning is utilized to effectively align heterogeneous cross-modal spatial and feature information. Finally, an attention mechanism is exploited to generate the fused representations by integrating spatial and feature information.

##### Construct feature neighbor graph and encode features

In a complex tissue sample, cells having the same cell types/states might not be in spatial adjacency with one another; in fact, they could be located quite far apart. To preserve both spatial adjacent information and the unique characteristics of non-adjacent cells with similar cell types/states, we propose to construct both the spatial graph 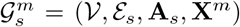 and the feature graph 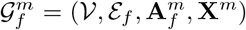 for the *m* ∈ {1, 2}-th modality. For lower-quality modality 1, the enhanced profiles 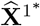 generated by denoising phase are adopted as feature matrix **X** . Besides, by applying KNN algorithm, spatial adjacency matrix **A**_*s*_ ∈ ℝ^*N* ×*N*^ is calculated according to the spatial coordinates 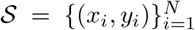, and the adjacency matrix of the feature graph 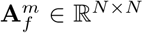 is computed according to the cosine similarity of expression feature matrix **X**^*m*^. By default, *k* set as 3 and 20 for spatial graph and feature graph.

Next, for each modality, we employed a parameter-sharing GCN to extract information separately from the spatial graph and the feature graph. By applying a shared-parameter GCN, we ensure that spatial and feature-based embeddings are aligned in a common representation space, which facilitates better integration of multi-modal information. The spatial graph captures local tissue organization by encoding physical spot proximity, whereas the feature graph preserves molecular similarity, linking spots that share similar expression profiles despite being spatially distant. This dual-graph structure allows CANDIES to effectively learn both spatially and phenotypically relevant patterns, ensuring that biologically meaningful relationships are retained. Specifically, the representations of cells are updated by iteratively aggregating information from their neighboring nodes in each graph, the cell’s embeddings in the *l*-th layer for the spatial graph 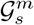 and the feature graph 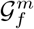 can be formalized as:

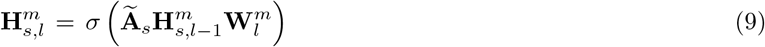

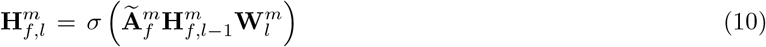

where the initial feature vectors of two graphs for *N* spots in GCN are both **X**^*m*^, i.e., 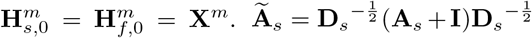 is the symmetrically normalized spatial adjacency matrix,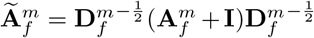 is the normalized feature similarity matrix. 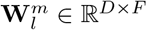 is the shared weight matrix in the *l*-th layer of GCN. *σ* denotes the ReLU activation function. After *L* layers, the final spatial and feature embedding outputs of GCN in Eq. (9) are 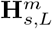 and 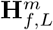.

##### Contrastive loss for cross-modality alignment

Contrastive learning [54] is a self-supervised learning technique that aims to learn useful representations by maximizing the similarity between positive pairs (similar samples) and minimizing the similarity between negative pairs (dissimilar samples). In spatial multi-omics data, each spot comprises two modalities, and *m*-th modality has spatial and feature embeddings (i.e., 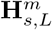 and 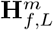) calculated by Eq. (9). In order to obtain effective integrated spot embeddings, a contrastive loss is employed to pull the spatial or feature embeddings of the same spot across the two modalities closer while pushing apart that of different spots. By applying contrastive learning, CANDIES learns the shared latent space that is invariant to the modality, allowing better alignment among different modalities. For spatial embeddings 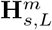, the formulation is:

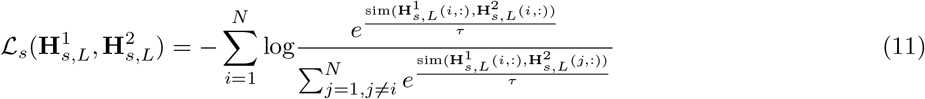

where sim(,) represents the similarity measurement function, *τ* is a temperature parameter, and 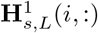 denotes the spatial embedding of the *i*-th cell in the 1st modality. Similarly, for feature embeddings 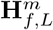, we can also obtain the formulation:

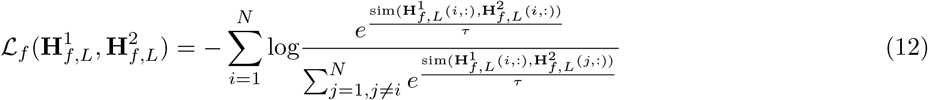

The overall contrastive loss is :

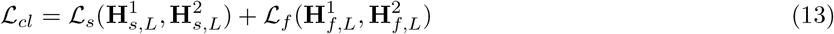

##### Integrating spatial and feature embeddings by attention

To further integrate the information from both modalities, we compute the mean of the spatial embeddings and feature embeddings across modalities for each spot. Specifically, the aggregated spatial embeddings **H**_*s*_ and feature embeddings **H**_*f*_ are derived as follows:

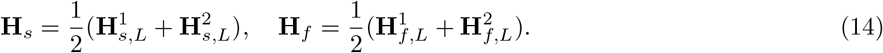

These aggregated representations capture shared biological structures and spatial relationships across different modalities. Following this, to effectively combine the aggregated spatial embeddings **H**_*s*_ and feature embeddings **H**_*f*_ into a unified representation **H**, we introduce a learnable attention mechanism that dynamically adjusts the contributions of two types of embeddings based on their relative importance for each cell:

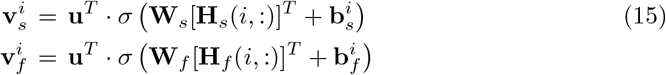

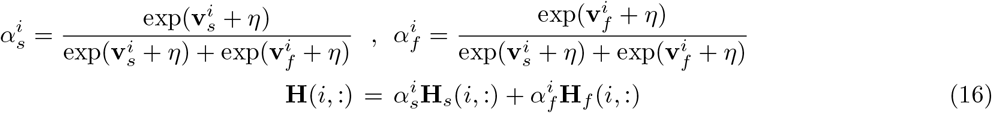

where 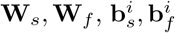 and **u** are trainable parameters, and 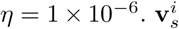 and 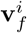 are the attention coefficient, while 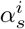 and 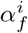 are the normalized attention scores, representing the importance of spatial or feature information to the representation of cell *i*.

##### Feature reconstruction loss

To ensure the fused representation **H** retains biologically meaningful information from spatial multi-omics, we introduce a feature reconstruction function 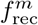 for each modality, which maps the fused representation back to the original feature space:

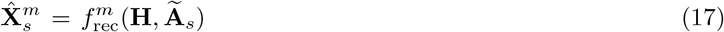

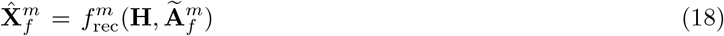

where 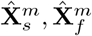 are the reconstructed feature of modality *m* using GCN on spatial graph and feature graph. The reconstruction loss is then defined as the mean squared error (MSE) between the original input features **X**^*m*^ and the reconstructed features:

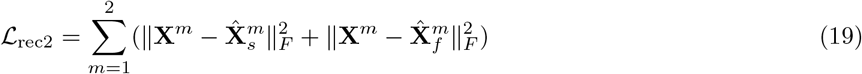

By incorporating the reconstruction loss, CANDIES maintains fidelity to the raw data while leveraging cross-attention to integrate spatial and feature information.

##### Integration model training

Combining the contrastive loss ℒ_*cl*_ in Eq. (13) and feature reconstruction loss ℒ_rec2_ in Eq. (19), the overall loss of integration phase in CANDIES is as follows:

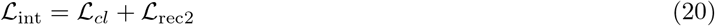

The final fused representation **H** by optimization Eq. (20) can be used in various downstream analysis, including domain identification, pseudo-spatiotemporal map construction, trait-domain association map generation.

#### CANDIES training details

As CANDIES follows a two-stage framework, we detailed the training process for each phase. In the denoising phase, the pre-trained graph auto-encoder was trained for 400 epochs, with the latent dimensionality fixed as 64 across all datasets. In the conditional diffusion model, CANDIES utilizes AdamW optimizer with a default learning rate of 0.001. To maintain consistency with traditional diffusion models, we partitioned the datasets into *k* folds based on the number of spots, where *k* was set as 3 by default. The conditional diffusion model was trained for a maximum of 1,000 epochs, with the number of diffusion steps as 800. In the integration phase, the optimizer was Adam, also with a default learning rate of 0.001. The maximum number of total training epochs in this phase was set as 300.

### Lower-quality modality determination in denoising phase

In the denoising phase, CANDIES aims to leverage the higher-quality modality to refine the lower-quality modality. Therefore, determining the relative quality between modalities is a crucial step. To achieve this, we established the following modality quality evaluation principles. During the denoising phase, we first pre-trained a dual graph auto-encoder to obtain low-dimensional embeddings for all spots across both modalities. We applied the Leiden algorithm on these embeddings to derive clustering results for each modality. Based on the clustering outcomes, we identified the lower-quality modality using several key metrics. When manual annotations or ground truth labels are available, we employed the Adjusted Rand Index (ARI), which is a widely used supervised metric, and selected the modality with the lower ARI as the lower-quality one. In the absence of manual annotations, we determined the lower-quality modality using four well-established unsupervised metrics: the Silhouette Index (SI), Davies-Bouldin Index (DBI), Moran’s I, and the Calinski-Harabasz Index (CHI). A modality with a lower DBI or higher SI, Moran’s I, and CHI is considered to have higher quality.

## Downstream analysis

### Pseudo-spatiotemporal map generation

To generate the pseudo Spatiotemporal Map (pSM) of the tissue, we computed the pSM value for each spot (or cell) by using *scanpy*.*tl*.*dpt* function implemented by scanpy package (v.1.10.3) [47]. We manually set the root spot to be the first spot of a given spot type. In spatial CITE-seq human skin data, MISAR-seq E18.5 mouse brain data, and spatial ATAC–RNA-seq mouse embryo data, the spot types selected to define root were dermis region, diencephalon_and_hindbrain region and forebrain region.

### Pseudo-time associated genes

For each gene, we calculated the pearson correlation coefficient between its expression across cells and the pseudo-spatiotemporal map (pSM) values derived from CANDIES. Genes exhibiting a significant correlation with a threshold of P ≤ 1e-10 were identified as pseudo-time-associated genes.

### Cluster-specific differentially expressed genes (DEGs)

For each cluster, we performed differential expressed genes analysis using the *scanpy*.*tl*.*rank genes groups* function from the scanpy package (v.1.10.3), and chose the Wilcoxon rank-sum test as the statistical method. A gene was classified as differentially expressed if its adjusted p-value (Q-value) was below the significance threshold of 0.05.

### GO enrichment analysis

GO (Gene Oncology) enrichment analysis were performed on pseudo-time-associated genes and cluster-specific DEGs using the *gseapy*.*enrichr* function implemented in the gseapy package (v.1.1.6)[55]. The analysis was conducted with the *GO_Biological_Process_2021* database as the reference, enabling the identification of enriched biological processes associated with the identified gene sets.

### Spatially resolved trait association mapping

We carried out gsMap analysis using the CANDIES embeddings as input to identify trait–spot associations across three tissues, including the mouse brain, mouse embryo, and human lymph. In the gsMap analysis, we directly computed cosine similarity and GSS using the latent embeddings from CANDIES. gsMap assigns the GSS of each spot as annotations to SNPs within a window extending 50 kb upstream and 50 kb downstream of each gene’s transcribed region, along with SNP-to-gene maps established using epigenomic data[42]. Within the S-LDSC framework, we considered stratified LD scores with respect to both baseline annotations and GSS annotations, thus obtaining the GSS enrichment p-value conditional on the baseline annotations. The baseline annotations include: (1) a constant value of 1 for all SNPs; (2) binary values (0 or 1) to indicate whether SNPs are mapped to genes; and (3) binary values to indicate whether SNPs are mapped to other functional annotations, including coding, conserved, and regulatory regions (e.g., promoter, enhancer, histone marks). Finally, to quantify the significance of the association of a specific spatial region with a trait, gsMap utilizes the Cauchy combination test to aggregate the p-values of individual spots within that spatial region.

## Data and Baselines

### Data information

Detailed information of spatial multi-omics datasets used in this manuscript was listed in the Supplementary Table. S1.

### Baseline methods

For the denoising task, seven methods designed for spatial transcriptomic data are included for benchmarking:

- GAAEST: https://github.com/tqwang743/GAAEST-main
- SCAN-IT: https://github.com/zcang/SCAN-IT
- SpaGCN: https://github.com/jianhuupenn/SpaGCN
- SEDR: https://github.com/JinmiaoChenLab/SEDR
- SpaceFlow: https://github.com/hongleir/SpaceFlow
- STAGATE: https://github.com/zhanglabtools/STAGATE
- GraphST: https://github.com/JinmiaoChenLab/GraphST

For the integration task, seven methods designed for multi-omics data are included for benchmarking:

- MultiVI, totalVI: https://github.com/scverse/scvi-tools
- scMDC: https://github.com/xianglin226/scMDC
- SpatialGlue: https://github.com/JinmiaoChenLab/SpatialGlue
- PRESENT: https://github.com/lizhen18THU/PRESENT
- PRAGA: https://github.com/Xubin-s-Lab/PRAGA
- COSMOS: https://github.com/Lin-Xu-lab/COSMOS

Among these baselines, SEDR is specially designed for spatial transcriptomic data denoising. SpaGCN was performed solely based on spatial transcriptomic data, without incorporating histological image information. MultiVI and totalVI were implemented via the scvi-tools package (v.1.3.0). We followed the tutorials provided by each method to conduct the experiments.

### Evaluation metrics

In our experiments, the evaluation of domain identification performance is conducted using a comprehensive set of metrics. For datasets with available ground truth, we employed supervised clustering metrics, including Mutual Information (Mutual Info), Normalized Mutual Information (NMI), Adjusted Mutual Information (AMI), V-measure, Homogeneity, Completeness, Adjusted Rand Index (ARI) and Fowlkes-Mallows Index (FMI). These metrics are designed to quantify the agreement between the predicted clusters and the annotated labels, offering valuable insights into the accuracy and consistency of the clustering results. On the other hand, for datasets without annotated labels, we utilized unsupervised clustering evaluation metrics, such as the Silhouette Index (SI), Davies-Bouldin Index (DBI), Moran’s I, and Calinski-Harabasz Index (CHI). These metrics assess clustering quality by measuring intra-cluster cohesion, inter-cluster separation, spatial autocorrelation, and variance ratios, enabling a robust evaluation of clustering structures in the absence of annotated data. For the calculation of Moran’s I, we employed the esda package (v.2.7.0) [56]. For other metrics, we utilized functions from the scikit-learn package (v.1.5.2) [57]. The detailed introduction of metrics used in this paper are listed as follows:

#### Mutual Information (MI)

it is a fundamental concept in information theory that measures the amount of information shared between the predicted labels and the true labels, ranging from 0 to positive values. A higher MI value indicates a stronger relationship between the predicted and the ground truth, the formulation is:

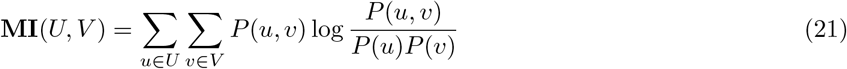

where *U* represents the set of ground truth, and *V* represents the set of predicted labels. *P* (*u*) and *P* (*v*) denote the marginal probabilities of the ground truth *u* and the predicted label *v, P* (*u, v*) represents the joint probabilities observing *u* and *v* simultaneously.

#### Normalized Mutual Information (NMI)

it is a normalized version of MI that adjusts for the bias introduced by different numbers of labels. It ensures that values remain between 0 and 1, where 1 indicates perfect clustering alignment, and 0 implies no correlation between predicted and ground truth labels. The NMI is defined as:

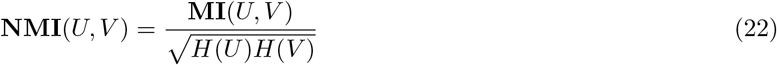

where 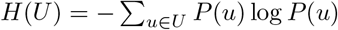 and 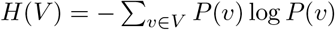 denote the entropy of the ground truth and predicted labels, respectively, measuring the uncertainty or variability in each label set.

#### Adjusted Mutual Information (AMI)

it is a variant of MI that corrects for randomness by adjusting the expected mutual information under a random clustering assumption. The higher the AMI value, the more similar the two sets of clusters are. AMI is calculated as follows:

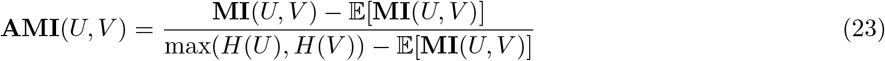

where 𝔼[**MI**(*U, V*)] represents the expected mutual information given a random labels assignment.

#### Homogeneity

Homogeneity measures whether each predicted cluster contains members of only a single ground truth category. The value of homogeneity ranges from 0 to 1, a higher homogeneity score means that each predicted cluster contains primarily samples from a single ground truth category. Homogeneity is defined as:

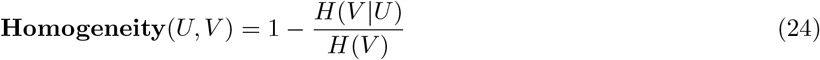

where 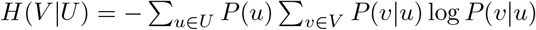 is the conditional entropy of predicted labels *V* given the true labels *U* .

#### Completeness

Completeness evaluates whether all members of a ground truth labels are assigned to the same predicted labels, a higher completeness score indicates that samples belonging to the same ground truth class are well grouped into the same predicted labels. Completeness is computed as:

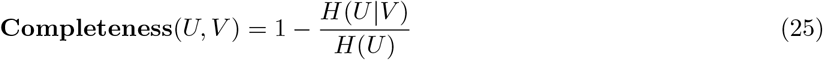

where 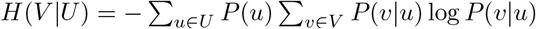 is the conditional entropy of true labels *U* given the predicted labels *V* .

#### V-measure

V-measure is a clustering evaluation metric that balances homogeneity and completeness to assess clustering quality. V-measure range from 0 to 1, where 1 indicates perfect clustering and 0 imply poor clustering, defined as:

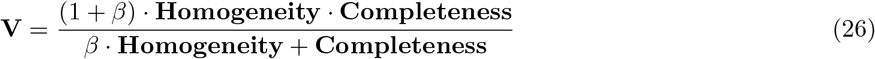

where *β* controls the weighting between homogeneity and completeness, typically set to 1 for an equal balance.

#### Adjusted Rand Index (ARI)

ARI is a clustering similarity measure that corrects the Rand Index (RI) for random chance. It evaluates how well the predicted clusters match the ground truth labels while considering random assignments. The ARI is computed as:

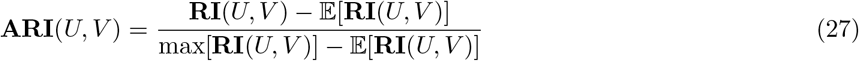

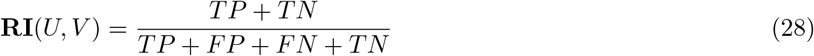

where *TP* is the number of true positives, *TN* is the number of true negatives, *FP* is the number of false positives, and *FN* is the number of false negatives.

#### Fowlkes-Mallows Index (FMI)

FMI measures the geometric mean of precision and recall between the true labels and predicted labels. FMI is defined as:

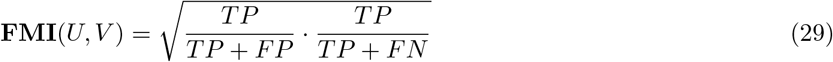

a higher FMI score indicates better clustering quality, with 1 being ideal and 0 indicating poor clustering performance.

#### Silhouette Index (SI)

SI measures the quality of clustering by evaluating both intra-labels cohesion and inter-labels separation. It is defined for each spot *i* and then averaged over all the samples. SI ranges from -1 to 1, where a high value indicates that the spot is well matched to its own labels and poorly matched to neighboring labels. SI is defined as:

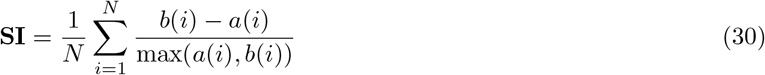

where *a*(*i*) is the average distance of sample *i* to other spots in the same label, and *b*(*i*) is the minimum average distance of *i* to spots in other labels.

#### Davies-Bouldin Index (DBI)

DBI evaluates clustering quality by measuring the average similarity between each label and its most similar label, based on the ratio of intra-label dispersion to inter-label separation. A lower DB score indicates better clustering, and it is defined as:

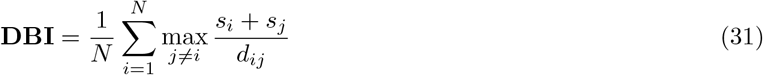

where *s*_*i*_ is the average intra-label distance for label *i*, and *d*_*ij*_ is the distance between the centroids of labels *i* and *j*.

#### Moran’s I

Moran’s I is a statistical measure used to evaluate spatial autocorrelation, determining whether similar values are clustered, dispersed, or randomly distributed across a spatial setting. It is widely employed in spatial clustering tasks to assess the degree of spatial dependence within the data. It is computed as

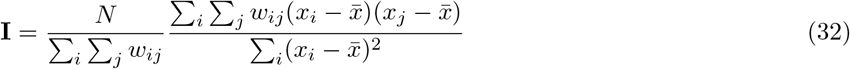

where *N* is the total number of observations, *x*_*i*_ and *x*_*j*_ are the observed values, 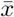 is the mean of *x, w*_*ij*_ represents the spatial weight between *i* and *j*. Moran’s I ranges from -1 to 1, where values close to 1 indicate positive spatial autocorrelation (clustering of similar values), values close to -1 suggest negative spatial autocorrelation (dispersion of dissimilar values), and values near 0 imply a random spatial distribution.

#### Calinski-Harabasz Index (CHI)

CHI also known as the Variance Ratio Criterion (VRC), evaluates clustering quality defined as the ratio of the between-cluster separation (*B*) to the within-cluster dispersion (*W*), normalized by their number of freedom degrees. A higher score indicates better-defined clusters. CHI is given by

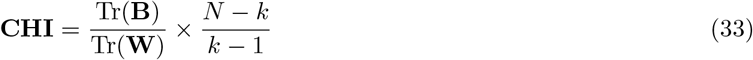

where **B** (between-cluster scatter matrix) is the weighted sum of squared Euclidean distances between each cluster centroid and the overall data centroid: 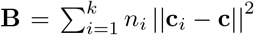 is the number of samples in cluster *C*_*i*_, **c**_*i*_ is the centroid of *C*_*i*_, and **c** is the overall centroid of the data. **B** measures how well the clusters are separated from each other (the higher, the better). **W** (within-cluster sum of squares) is the sum of squared Euclidean distances between the data points **x** and their respective cluster centroids: 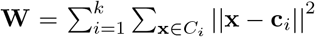. **W** measures the compactness or cohesiveness of the clusters (the smaller, the better). Furthermore, Tr() is the trace function, *N* is the total number of samples, and *k* is the number of clusters.

## Data availability

The original code to simulate the spatial multi-omics data is from https://github.com/willtownes/nsf-paper. The MISAR-seq mouse brain data is obtained from https://www.biosino.org/node/project/detail/OEP003285. The spatial CITE-seq sequencing data from human skin biopsy tissue is available in the Gene Expression Omnibus (GSE213264), and the high-resolution microscope image of the tissue is available at https://figshare.com/articles/figure/figures/20723680. The spatial ATAC–RNA-seq data from mouse embryo is available in the UCSC Cell and Genome (https://cells.ucsc.edu/?ds=brain-spatial-omics), and is also in the AtlasX-plore (https://web.atlasxomics.com/visualization/Fan). The in-house human lymph node data is obtained from SpatialGlue and is available at https://zenodo.org/records/7879713. All the data used in the manuscript have been uploaded to Zenodo and are freely available at https://zenodo.org/records/15070780.

## Code availability

The implementation of CANDIES is accessible at https://github.com/YeLiu-Lab/CANDIES. The tutorials for analyzing the datasets used in this paper are available at https://github.com/YeLiu-Lab/CANDIES/tree/main/CANDIES/tutorial.

## References

[1] Katy Vandereyken et al. “Methods and applications for single-cell and spatial multi-omics”. In: Nature Reviews Genetics 24.8 (2023), pp. 494–515.

[2] Yang Liu et al. “High-spatial-resolution multi-omics sequencing via deterministic barcoding in tissue”. In: Cell 183.6 (2020), pp. 1665–1681.

[3] Yang Liu et al. “High-plex protein and whole transcriptome co-mapping at cellular resolution with spatial CITE-seq”. In: Nature Biotechnology 41.10 (2023), pp. 1405–1409.

[4] Sha Liao et al. “Integrated spatial transcriptomic and proteomic analysis of fresh frozen tissue based on stereo-seq”. In: bioRxiv (2023), pp. 2023–04.

[5] Shanshan He et al. “High-plex imaging of RNA and proteins at subcellular resolution in fixed tissue by spatial molecular imaging”. In: Nature Biotechnology 40.12 (2022), pp. 1794–1806.

[6] Di Zhang et al. “Spatial epigenome–transcriptome co-profiling of mammalian tissues”. In: Nature 616.7955 (2023), pp. 113–122.

[7] Fuqing Jiang et al. “Simultaneous profiling of spatial gene expression and chromatin accessibility during mouse brain development”. In: Nature Methods 20.7 (2023), pp. 1048–1057.

[8] Paul Kiessling and Christoph Kuppe. “Spatial multi-omics: novel tools to study the complexity of cardiovascular diseases”. In: Genome Medicine 16.1 (2024), p. 14.

[9] Patrik L Ståhl et al. “Visualization and analysis of gene expression in tissue sections by spatial transcriptomics”. In: Science 353.6294 (2016), pp. 78–82.

[10] Linhua Wang, Mirjana Maletic-Savatic, and Zhandong Liu. “Region-specific denoising identifies spatial coexpression patterns and intra-tissue heterogeneity in spatially resolved transcriptomics data”. In: Nature Communications 13.1 (2022), p. 6912.

[11] Ruochen Jiang et al. “Statistics or biology: the zero-inflation controversy about scRNA-seq data”. In: Genome biology 23.1 (2022), p. 31.

[12] Kyle Coleman, Amelia Schroeder, and Mingyao Li. “Unlocking the power of spatial omics with AI”. In: nature methods 21.8 (2024), pp. 1378–1381.

[13] Yuhan Hao et al. “Integrated analysis of multimodal single-cell data”. In: Cell 184.13 (2021), pp. 3573–3587.

[14] Tal Ashuach et al. “MultiVI: deep generative model for the integration of multimodal data”. In: Nature Methods 20.8 (2023), pp. 1222–1231.

[15] Adam Gayoso et al. “Joint probabilistic modeling of single-cell multi-omic data with totalVI”. In: Nature methods 18.3 (2021), pp. 272–282.

[16] Xiuhui Yang et al. “scCross: a deep generative model for unifying single-cell multi-omics with seamless integration, cross-modal generation, and in silico exploration”. In: Genome Biology 25.1 (2024), p. 198.

[17] Pengfei Guo et al. “Multiplexed spatial mapping of chromatin features, transcriptome and proteins in tissues”. In: Nature Methods (2025), pp. 1–10.

[18] Jian Hu et al. “SpaGCN: Integrating gene expression, spatial location and histology to identify spatial domains and spatially variable genes by graph convolutional network”. In: Nature methods 18.11 (2021), pp. 1342–1351.

[19] Yahui Long et al. “Spatially informed clustering, integration, and deconvolution of spatial transcriptomics with GraphST”. In: Nature Communications 14.1 (2023), p. 1155.

[20] Hang Xu et al. “Unsupervised spatially embedded deep representation of spatial transcriptomics”. In: Genome Medicine 16.1 (2024), p. 12.

[21] Ruben Dries et al. “Giotto: a toolbox for integrative analysis and visualization of spatial expression data”. In: Genome biology 22 (2021), pp. 1–31.

[22] Lambda Moses and Lior Pachter. “Museum of spatial transcriptomics”. In: Nature methods 19.5 (2022), pp. 534–546.

[23] Luyi Tian, Fei Chen, and Evan Z Macosko. “The expanding vistas of spatial transcriptomics”. In: Nature Biotechnology 41.6 (2023), pp. 773–782.

[24] Emma Lundberg and Georg HH Borner. “Spatial proteomics: a powerful discovery tool for cell biology”. In: Nature Reviews Molecular Cell Biology 20.5 (2019), pp. 285–302.

[25] Yahui Long et al. “Deciphering spatial domains from spatial multi-omics with SpatialGlue”. In: Nature Methods (2024), pp. 1–10.

[26] Zhen Li et al. “Cross-modality representation and multi-sample integration of spatially resolved omics data”. In: bioRxiv (2024), pp. 2024–06.

[27] Yuansheng Zhou et al. “Cooperative integration of spatially resolved multi-omics data with COSMOS”. In: Nature Communications 16.1 (2025), p. 27.

[28] Xinlei Huang et al. “PRAGA: prototype-aware graph adaptive aggregation for spatial multi-modal omics analysis”. In: arXiv preprint 2409.12728 (2024).

[29] William Peebles and Saining Xie. “Scalable diffusion models with transformers”. In: Proceedings of the IEEE/CVF International Conference on Computer Vision. 2023, pp. 4195–4205.

[30] Thomas N Kipf and Max Welling. “Semi-supervised classification with graph convolutional networks”. In: arXiv preprint 1609.02907 (2016).

[31] Tianqi Wang et al. “Graph attention automatic encoder based on contrastive learning for domain recognition of spatial transcriptomics”. In: Communications Biology 7.1 (2024), p. 1351.

[32] Zixuan Cang et al. “SCAN-IT: Domain segmentation of spatial transcriptomics images by graph neural network”. In: BMVC: proceedings of the british machine vision conference. british machine vision conference. Vol. 32. NIH Public Access. 2021.

[33] Honglei Ren et al. “Identifying multicellular spatiotemporal organization of cells with SpaceFlow”. In: Nature communications 13.1 (2022), p. 4076.

[34] Kangning Dong and Shihua Zhang. “Deciphering spatial domains from spatially resolved transcriptomics with an adaptive graph attention auto-encoder”. In: Nature communications 13.1 (2022), p. 1739.

[35] Vincent A Traag, Ludo Waltman, and Nees Jan Van Eck. “From Louvain to Leiden: guaranteeing well-connected communities”. In: Scientific reports 9.1 (2019), pp. 1–12.

[36] Laleh Haghverdi et al. “Diffusion pseudotime robustly reconstructs lineage branching”. In: Nature methods 13.10 (2016), pp. 845–848.

[37] Wolfram Sterry. Kurzlehrbuch Dermatologie. Georg Thieme Verlag, 2018.

[38] Xiang Lin et al. “Clustering of single-cell multi-omics data with a multimodal deep learning method”. In: Nature communications 13.1 (2022), p. 7705.

[39] JAMES Y LI et al. “Patterning and compartment formation in the diencephalon”. In: Frontiers in neuroscience 6 (2012), p. 23290.

[40] Qiwen Li et al. “Insights into skeletal stem cells”. In: Bone research 10.1 (2022), p. 61.

[41] Luciano Custo Greig et al. “Molecular logic of neocortical projection neuron specification, development and diversity”. In: Nature Reviews Neuroscience 14.11 (2013), pp. 755–769.

[42] Liyang Song et al. “Spatially resolved mapping of cells associated with human complex traits”. In: Nature (2025), pp. 1–10.

[43] Yitao Yang et al. “STAIG: Spatial transcriptomics analysis via image-aided graph contrastive learning for domain exploration and alignment-free integration”. In: Nature Communications 16.1 (2025), p. 1067.

[44] Bohan Li et al. “Tissue characterization at an enhanced resolution across spatial omics platforms with deep generative model”. In: Nature Communications 15.1 (2024), p. 6541.

[45] F William Townes and Barbara E Engelhardt. “Nonnegative spatial factorization applied to spatial genomics”. In: Nature methods 20.2 (2023), pp. 229–238.

[46] Tim Stuart et al. “Comprehensive integration of single-cell data”. In: cell 177.7 (2019), pp. 1888–1902.

[47] F Alexander Wolf, Philipp Angerer, and Fabian J Theis. “SCANPY: large-scale single-cell gene expression data analysis”. In: Genome biology 19 (2018), pp. 1–5.

[48] Matthew P Mulé, Andrew J Martins, and John S Tsang. “Normalizing and denoising protein expression data from droplet-based single cell profiling”. In: Nature communications 13.1 (2022), p. 2099.

[49] Ricard Argelaguet et al. “Computational principles and challenges in single-cell data integration”. In: Nature biotechnology 39.10 (2021), pp. 1202–1215.

[50] Xiaoyang Chen et al. “Cell type annotation of single-cell chromatin accessibility data via supervised Bayesian embedding”. In: Nature Machine Intelligence 4.2 (2022), pp. 116–126.

[51] Gökcen Eraslan et al. “Single-cell RNA-seq denoising using a deep count autoencoder”. In: Nature communications 10.1 (2019), p. 390.

[52] Jonathan Ho, Ajay Jain, and Pieter Abbeel. “Denoising diffusion probabilistic models”. In: Advances in neural information processing systems 33 (2020), pp. 6840–6851.

[53] Robin Rombach et al. “High-resolution image synthesis with latent diffusion models”. In: Proceedings of the IEEE/CVF conference on computer vision and pattern recognition. 2022, pp. 10684–10695.

[54] Ting Chen et al. “A simple framework for contrastive learning of visual representations”. In: International conference on machine learning. PMLR. 2020, pp. 1597–1607.

[55] Zhuoqing Fang, Xinyuan Liu, and Gary Peltz. “GSEApy: a comprehensive package for performing gene set enrichment analysis in Python”. In: Bioinformatics 39.1 (2023), btac757.

[56] Sergio J Rey and Luc Anselin. “PySAL: A Python library of spatial analytical methods”. In: Handbook of applied spatial analysis: Software tools, methods and applications. Springer, 2009, pp. 175–193.

[57] Fabian Pedregosa et al. “Scikit-learn: Machine learning in Python”. In: the Journal of machine Learning research 12 (2011), pp. 2825–2830.

