## Supplemental_note for "Cross-modal Denoising and Integration of Spatial Multi-omics data with CANDIES"

**Table S1**: Experimental datasets used in the manuscript.

| Dataset name | Platform (omics) | High-quality modality | Size (spots × genes/proteins/peaks) |
|---|---|---|---|
| Synthetic data | - (RNA-protein) | protein | 1,296 × 800 / 100 |
| Human Skin | Spatial CITE-seq (RNA-protein) | protein | 1,691 × 15,486 / 283 |
| E18.5 Mouse Brain | MISAR-seq (RNA-ATAC) | ATAC | 2,129 × 32,285 / 161,461 |
| E15.5 Mouse Brain | MISAR-seq (RNA-ATAC) | ATAC | 1,949 × 32,285 / 191,034 |
| E13 Mouse Embryo | Spatial ATAC–RNA-seq (RNA-ATAC) | RNA | 2,186 × 20,900 / 87,173 |
| Human Lymph Node | 10x Genomics Visium (RNA-protein) | RNA | 3,484 × 18,085 / 31 |

**Table S2**: Applicability of integration methods for different data types.

| | totalVI | MultiVI | scMDC | SpatialGlue | PRESENT | PRAGA | COSMOS | CANDIES |
|---|---|---|---|---|---|---|---|---|
| RNA & protein | ✓ | ✓ | ✓ | ✓ | ✓ | ✓ | ✓ | ✓ |
| RNA & ATAC | × | ✓ | ✓ | ✓ | ✓ | × | ✓ | ✓ |

**Table S3**: Detailed parameter setting of simulated data generation with varying standard deviation.

| **Modality 1(RNA)** | | | | | | **Modality 2(Protein)** | | | | |
|---|---|---|---|---|---|---|---|---|---|---|
| ZINB | | | Gaussian noise | | Dimension | NB | | Gaussian noise | | Dimension |
| pi | bkg_mean | expr_mean | mean | std | | bkg_mean | expr_mean | mean | std | |
| 0.3 | 0.5 | 1.5 | 2 | 2.0 | 800 | 2 | 3 | 2 | 1 | 100 |
| 0.3 | 0.5 | 1.5 | 2 | 2.5 | 800 | 2 | 3 | 2 | 1 | 100 |
| 0.3 | 0.5 | 1.5 | 2 | 3.0 | 800 | 2 | 3 | 2 | 1 | 100 |
| 0.3 | 0.5 | 1.5 | 2 | 3.5 | 800 | 2 | 3 | 2 | 1 | 100 |
| 0.3 | 0.5 | 1.5 | 2 | 4.0 | 800 | 2 | 3 | 2 | 1 | 100 |
| 0.3 | 0.5 | 1.5 | 2 | 4.5 | 800 | 2 | 3 | 2 | 1 | 100 |
| 0.3 | 0.5 | 1.5 | 2 | 5.0 | 800 | 2 | 3 | 2 | 1 | 100 |

**Table S4**: Detailed parameter setting of simulated data generation with varying dropout rate.

| **Modality 1(RNA)** | | | | | | **Modality 2(Protein)** | | | | |
|---|---|---|---|---|---|---|---|---|---|---|
| ZINB | | | Gaussian | | Dimension | NB | | Gaussian noise | | Dimension |
| pi | bkg_mean | expr_mean | mean | std | | bkg_mean | expr_mean | mean | std | |
| 0.2 | 0.5 | 1.5 | 2 | 1 | 800 | 2 | 3 | 2 | 1 | 100 |
| 0.3 | 0.5 | 1.5 | 2 | 1 | 800 | 2 | 3 | 2 | 1 | 100 |
| 0.4 | 0.5 | 1.5 | 2 | 1 | 800 | 2 | 3 | 2 | 1 | 100 |
| 0.5 | 0.5 | 1.5 | 2 | 1 | 800 | 2 | 3 | 2 | 1 | 100 |

**Table S5**: 32 complex traits used for downstream analysis

| Complex Trait | Abbreviation | Category | Link |
|---|---|---|---|
| Intelligence Quotient | IQ | Psychiatric | https://www.nature.com/articles/s41588-018-0152-6 |
| Height | Height | Anthropometric | https://www.nature.com/articles/s41586-022-05275-y |
| Schizophrenia | SCZ | Psychiatric | https://www.nature.com/articles/s41586-022-04434-5 |
| Major Depressive Disorder | MDD | Psychiatric | https://www.nature.com/articles/s41593-018-0326-7 |
| Mean Corpuscular Hemoglobin Concentration | MCHC | Haemotological | https://www.nature.com/articles/s41588-018-0047-6 |
| Mean Corpuscular Hemoglobin | MCH | Haemotological | https://www.nature.com/articles/s41588-018-0047-6 |
| Body Mass Index | BMI | Anthropometric | https://www.nature.com/articles/s41588-018-0144-6 |
| Mean Corpuscular Volume | MCV | Haemotological | https://www.nature.com/articles/s41588-018-0047-6 |
| Basophils | Baso | Haemotological | https://www.cell.com/cell/fulltext/S0092-8674(20)30999-5 |
| Monocytes | Mono | Haemotological | https://www.cell.com/cell/fulltext/S0092-8674(20)30999-5 |
| Neutrophils | Neutro | Haemotological | https://www.cell.com/cell/fulltext/S0092-8674(20)30999-5 |
| Eosinophils | Eosino | Haemotological | https://www.cell.com/cell/fulltext/S0092-8674(20)30999-5 |
| Platelets | Plt | Haemotological | https://www.cell.com/cell/fulltext/S0092-8674(20)30999-5 |
| Red Blood Cells | RBC | Haemotological | https://www.cell.com/cell/fulltext/S0092-8674(20)30999-5 |
| Hemoglobin | Hb | Haemotological | https://www.cell.com/cell/fulltext/S0092-8674(20)30999-5 |
| Lymphocytes | Lym | Haemotological | https://www.cell.com/cell/fulltext/S0092-8674(20)30999-5 |
| White Blood Cells | WBC | Haemotological | https://www.cell.com/cell/fulltext/S0092-8674(20)30999-5 |
| High-Density Lipoprotein | HDL | Metabolism | https://csg.sph.umich.edu/willer/public/glgc-lipids2021/ |
| Low-Density Lipoprotein | LDL | Metabolism | https://csg.sph.umich.edu/willer/public/glgc-lipids2021/ |
| Total Cholesterol | TC | Metabolism | https://csg.sph.umich.edu/willer/public/glgc-lipids2021/ |
| Log-transformed Triglycerides | logTG | Metabolism | https://csg.sph.umich.edu/willer/public/glgc-lipids2021/ |
| Attention Deficit Hyperactivity Disorder | ADHD | Psychiatric | https://pgc.unc.edu/for-researchers/download-results/ |
| Alzheimer's Disease | ALZ | Psychiatric | https://pgc.unc.edu/for-researchers/download-results/ |
| Anxiety Disorder | ANX | Psychiatric | https://pgc.unc.edu/for-researchers/download-results/ |
| Autism Spectrum Disorder | ASD | Psychiatric | https://pgc.unc.edu/for-researchers/download-results/ |
| Bipolar Disorder | BIP | Psychiatric | https://pgc.unc.edu/for-researchers/download-results/ |
| Obsessive-Compulsive Disorder / Tourette Syndrome | OCDTS | Psychiatric | https://pgc.unc.edu/for-researchers/download-results/ |
| Post-Traumatic Stress Disorder | PTSD | Psychiatric | https://pgc.unc.edu/for-researchers/download-results/ |
| Education attainment | EA | Behaviour | https://www.nature.com/articles/s41588-018-0147-3 |
| Drinks per week | DPW | Behaviour | https://www.nature.com/articles/s41588-018-0307-5 |
| Smoking Cessation | SmokingC | Behaviour | https://www.nature.com/articles/s41588-018-0307-5 |
| Smoking Initiation | SmokingI | Behaviour | https://www.nature.com/articles/s41588-018-0307-5 |

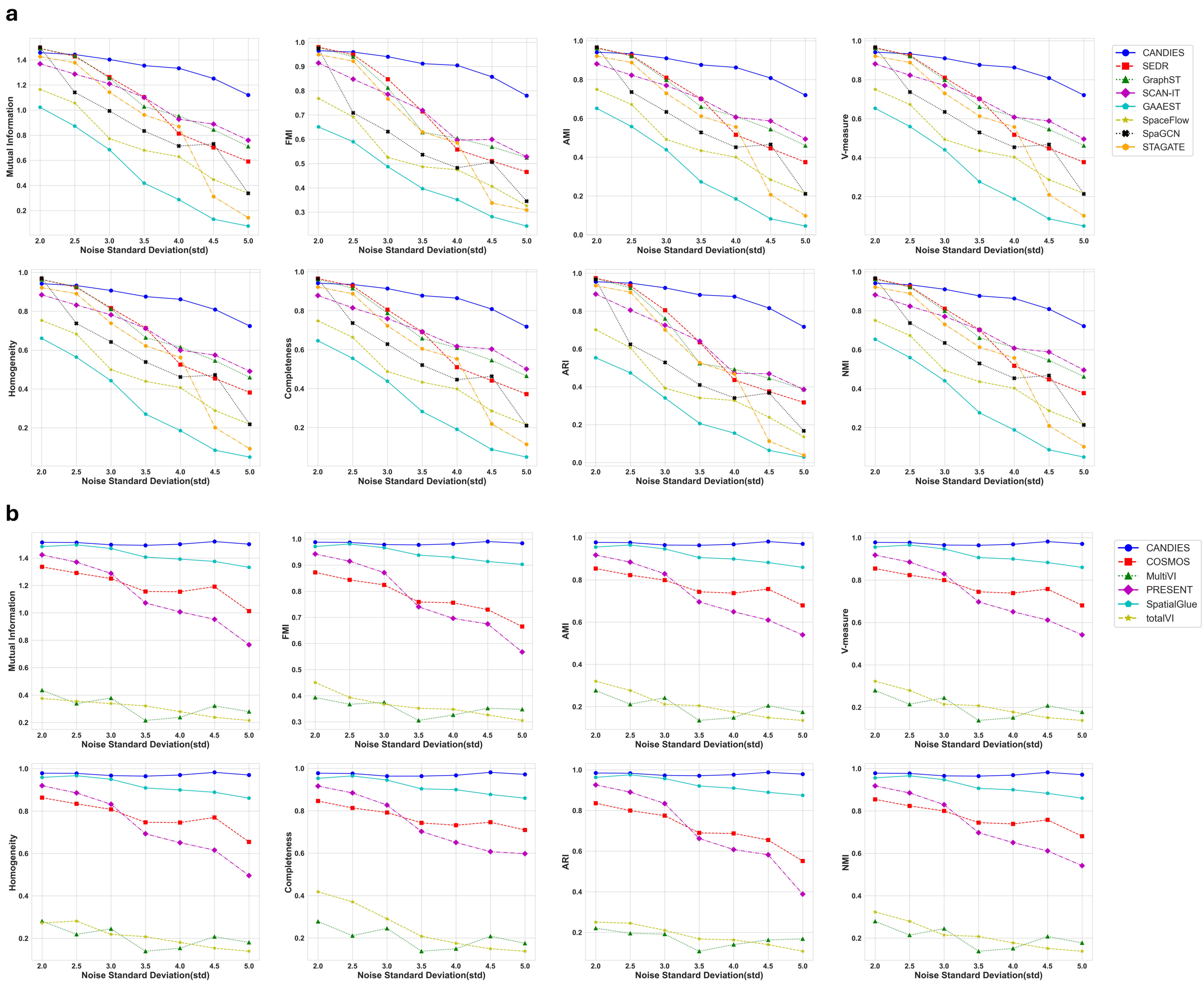


**Fig. S1**: **Quantitative analysis of simulated data generated with varying standard deviations.** **a**, Spatial transcriptome denoising results across eight supervised evaluation metrics. **b**, Multi-omics integration results across eight supervised evaluation metrics.

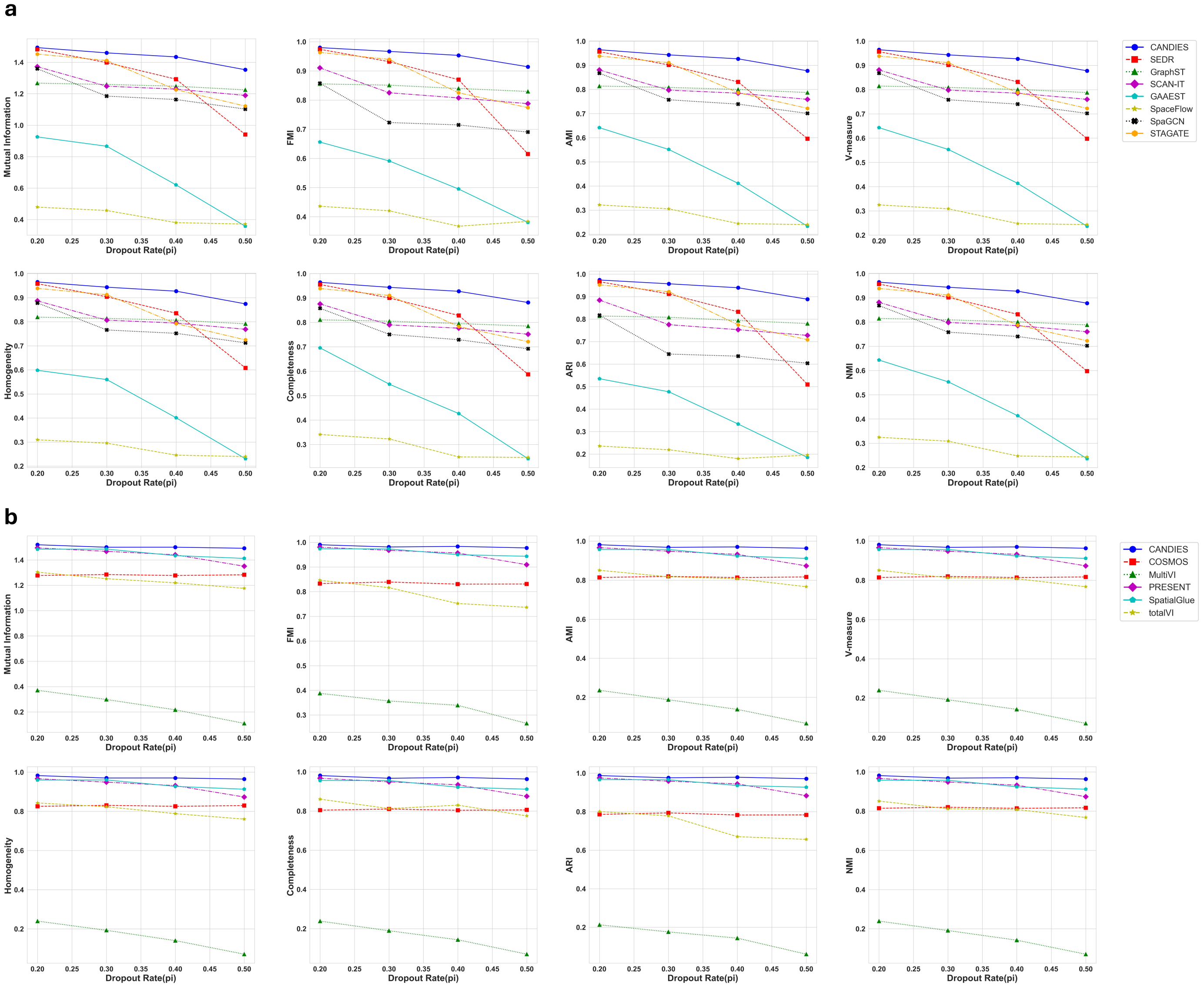


**Fig. S2**: **Quantitative analysis of simulated data generated with varying dropout rate. a**, Spatial transcriptome denoising results across eight supervised evaluation metrics. **b**, Multi-omics integration results across eight supervised evaluation metrics.

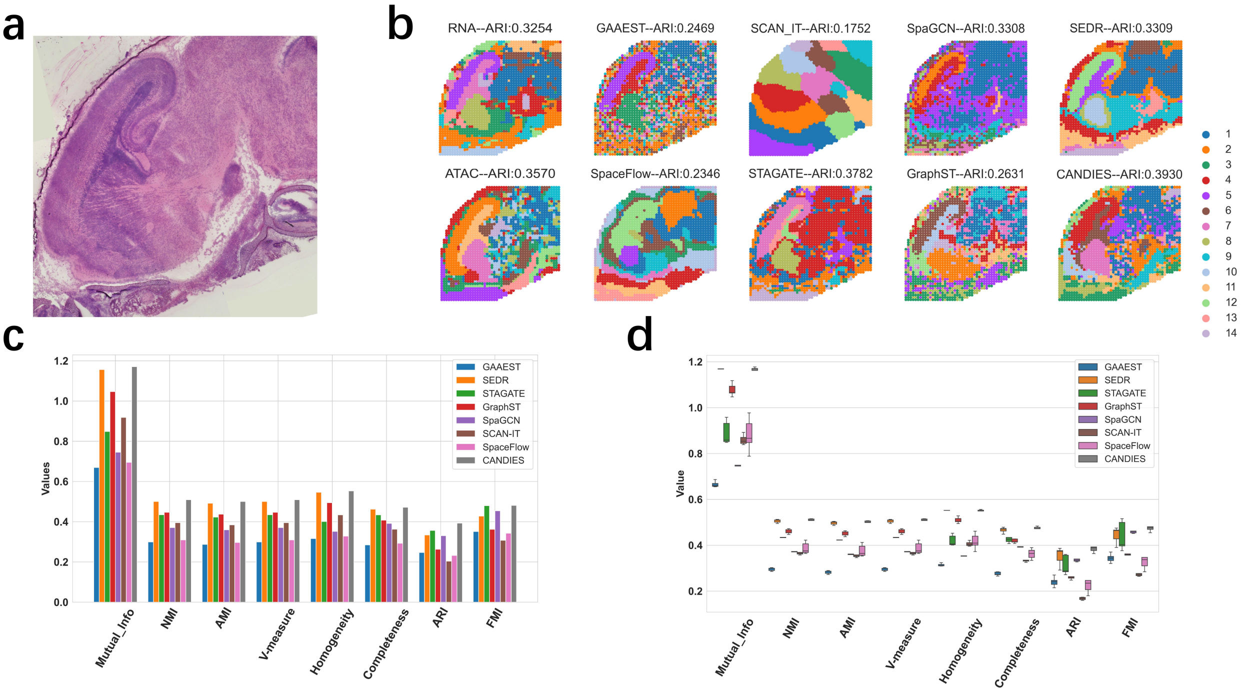


**Fig. S3**: **Denoising results for E18.5 mouse brain data. a**, The bright-field image of the E18.5 mouse brain tissue section. **b**, Spatial plots of the E18.5 mouse brain data only with RNA modality across eight representative spatial transcriptomics (ST) methods, the first column is identified by Leiden after encoding phase. **c**, Bar plots comparing the performance of eight methods across eight supervised metrics. **d**, Box plots comparing the performance of eight methods across eight supervised metrics.

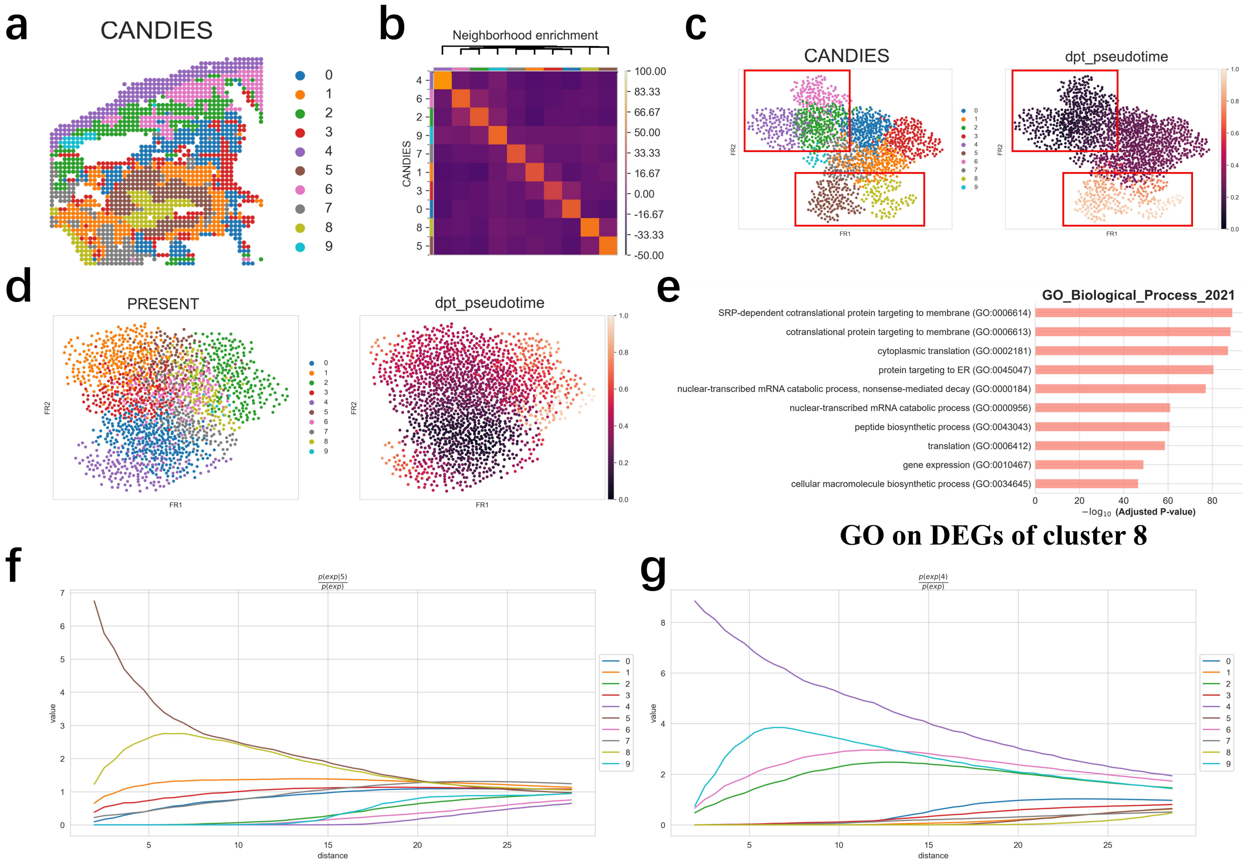


**Fig. S4**: **Additional results of the human skin data after integration.** **a**, Spatial plot of CANDIES integration clustering result. **b**, Similarity heatmap of clusters identified by CANDIES, illustrating the spatial proximity relationships between different clusters. **c**, The UMAP visualization of CANDIES' latent representations colored by clusters (left) and pSM values (right). **d**, The UMAP visualization of PRESENT's latent representations colored by clusters (left) and pSM values (right). **e**, Gene Ontology (GO) enrichment analysis on differentially expressed genes (DEGs) of cluster 8. **f**, Cluster co-occurrence score for cluster 5 at increasing distances. **g**, Cluster co-occurrence score for cluster 4 at increasing distances.

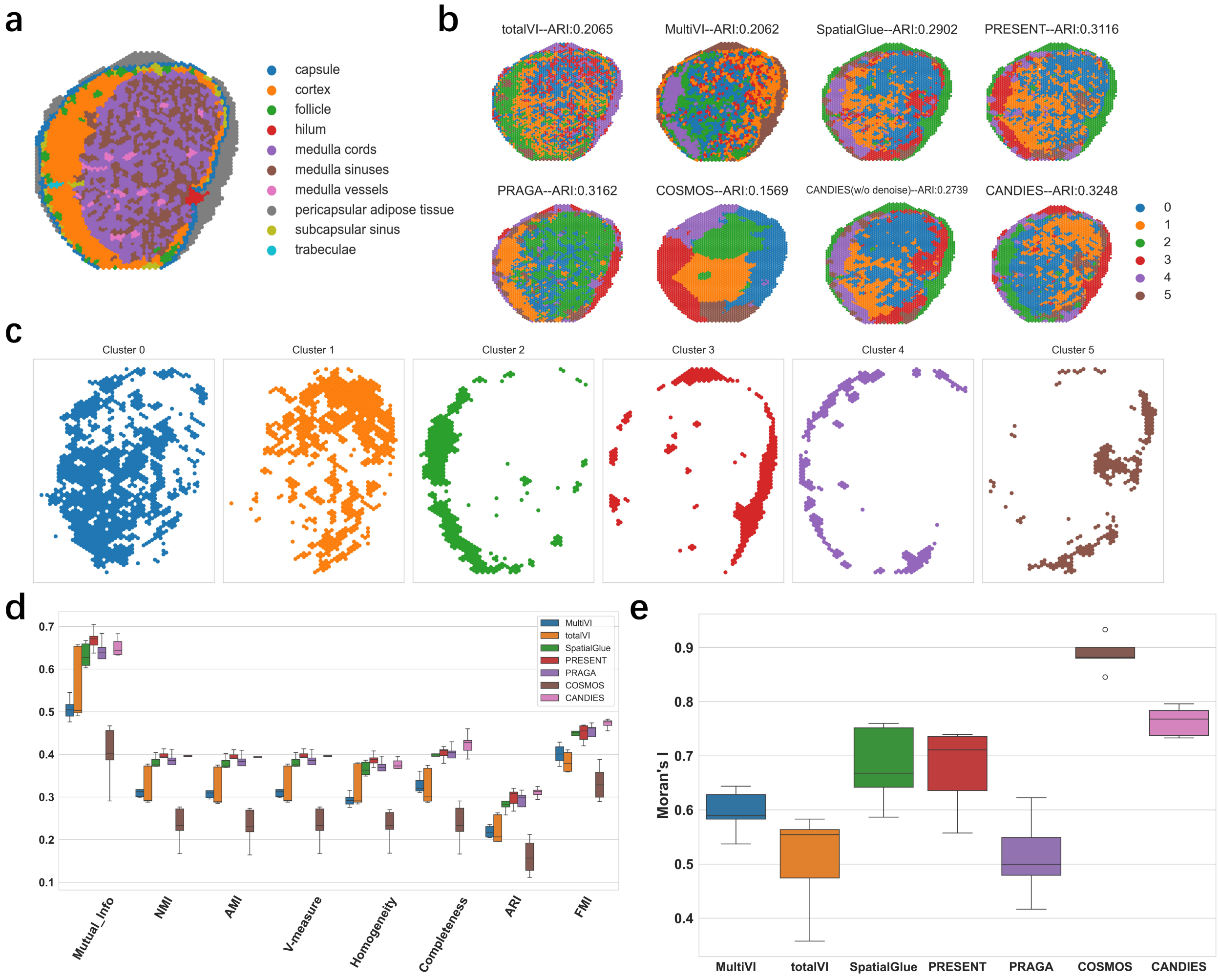


**Fig. S5**: **Integration results for human lymph node data. a**, Manual annotation of the human lymph node data. **b**, Spatial plots of the human lymph node data across seven single-cell and spatial multi-omics integration methods, CANDIES (w/o denoise) refers to the variant of CANDIES without the denoising phase. **c**, Separate spatial plots of all clusters identified by CANDIES in the human lymph node data. **d**, Box plots of the eight supervised metrics across the seven methods. **e**, Box plots of the Moran's I scores across the seven methods.

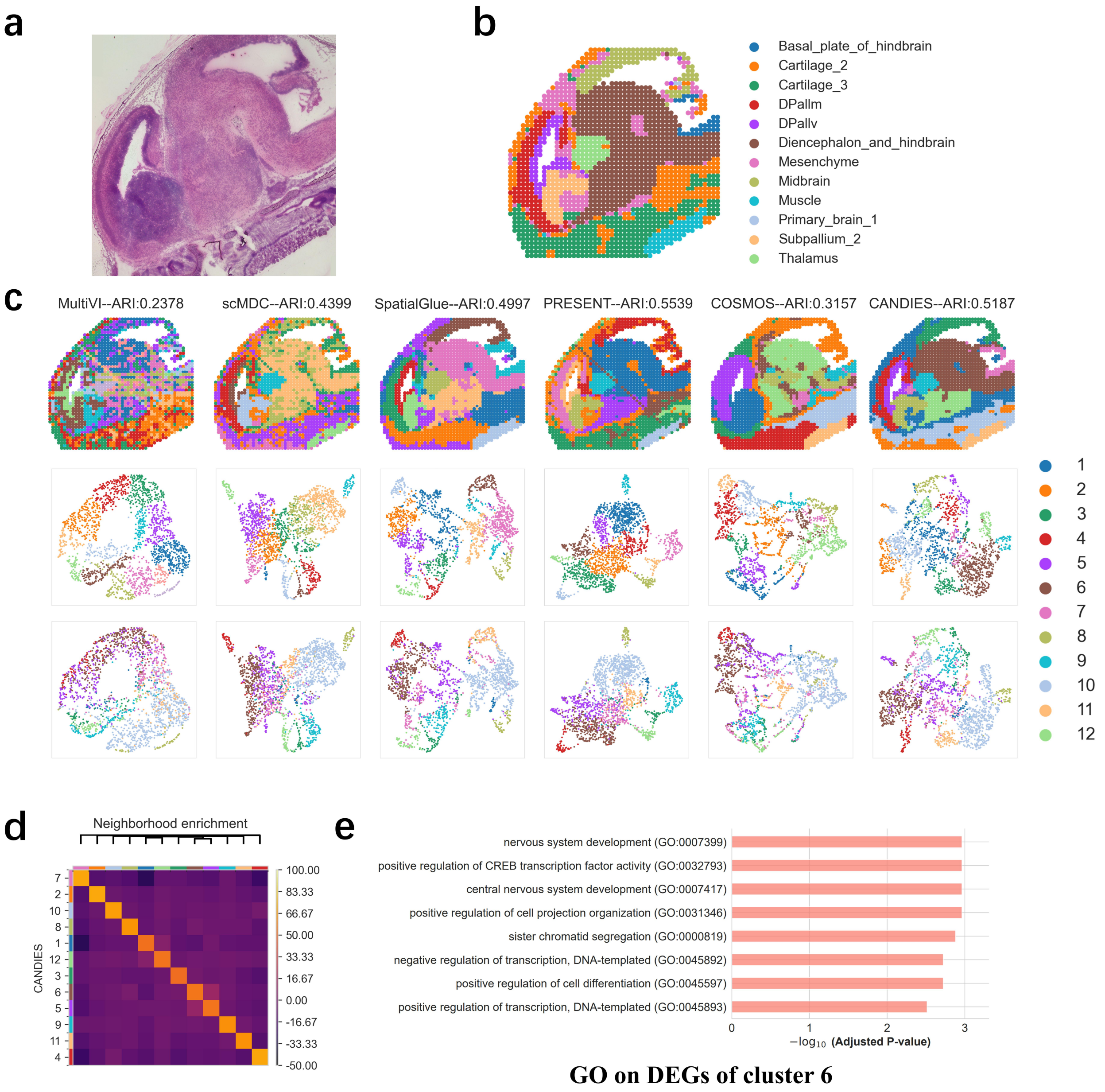


**Fig. S6**: **Integration results for E15.5 mouse brain data.** **a**, The bright-field image of the E15.5 mouse brain tissue section. **b**, Manual annotation of the E15.5 mouse brain data. **c**, Spatial plots and UMAP visualization of the E15.5 mouse brain data across six single-cell and spatial multi-omics integration methods, the second row is colored by clusters identified by CANDIES, and the third row is colored by manual annotations. **d**, Similarity heatmap of clusters identified by CANDIES, illustrating the spatial proximity relationships between different clusters. **e**, Gene Ontology (GO) enrichment analysis on differentially expressed genes (DEGs) of cluster 6.

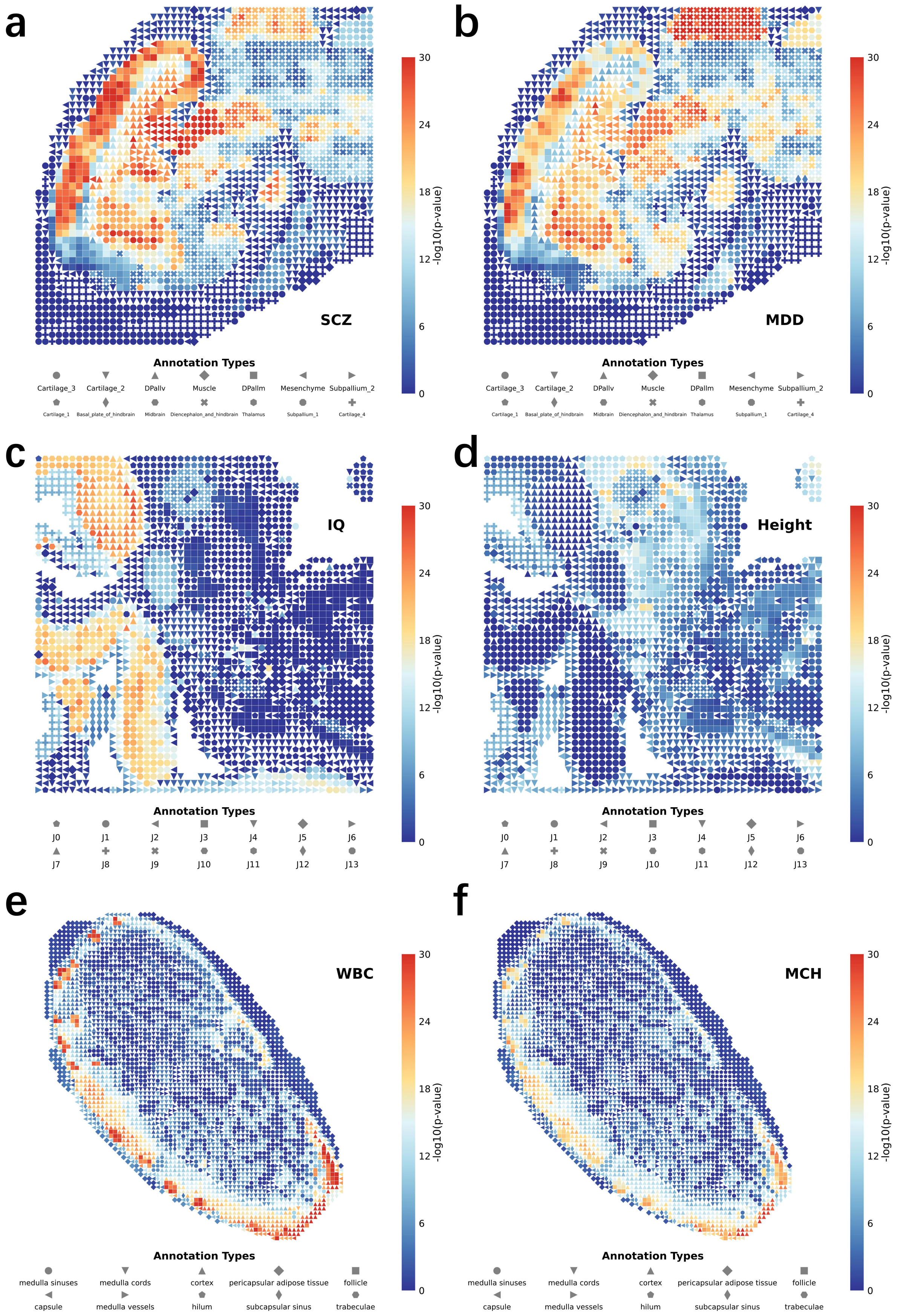


**Fig. S7**: **Trait-spot associations identified by using gsMap embeddings in three spatial omics datasets as input. a,b**, Two psychiatric traits (major depressive disorder and schizophrenia) highlighted in mouse brain, with spot annotation indicated by different point shapes and $\log_{10}$ $p$-values reflected by the color. **c,d**, Two representative traits (IQ and height) in mouse embryo, illustrating how developmental regions associate with cognition and anthropometric traits. **e,f**, Two hematological traits (white blood cells count and mean corpuscular hemoglobin) in human lymph, illustrating localized enrichment patterns.

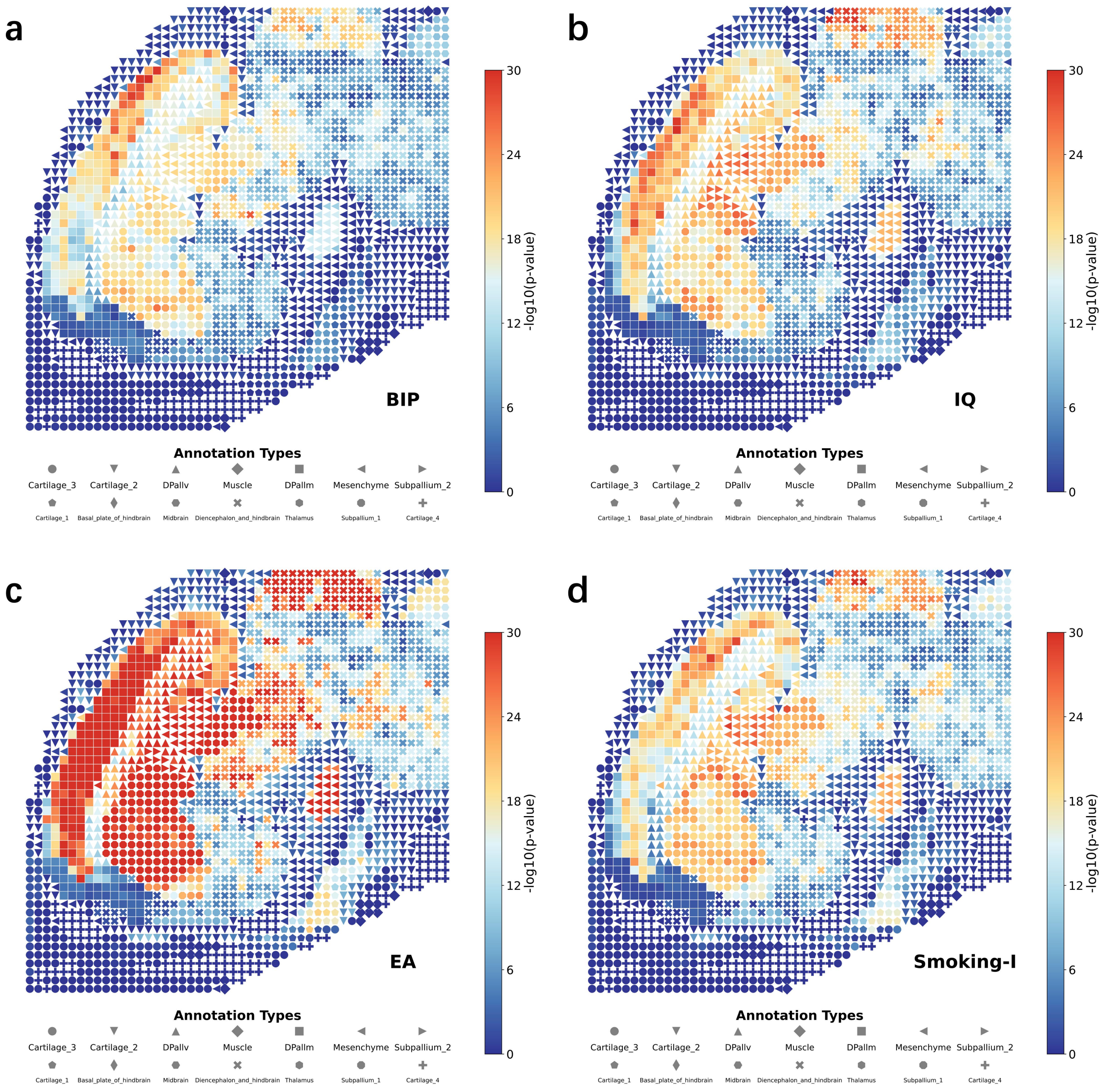


**Fig. S8**: Trait-spot associations for BIP, IQ, EA and Smoking-I traits identified by using CANDIES embeddings in mouse brain datasets as input.

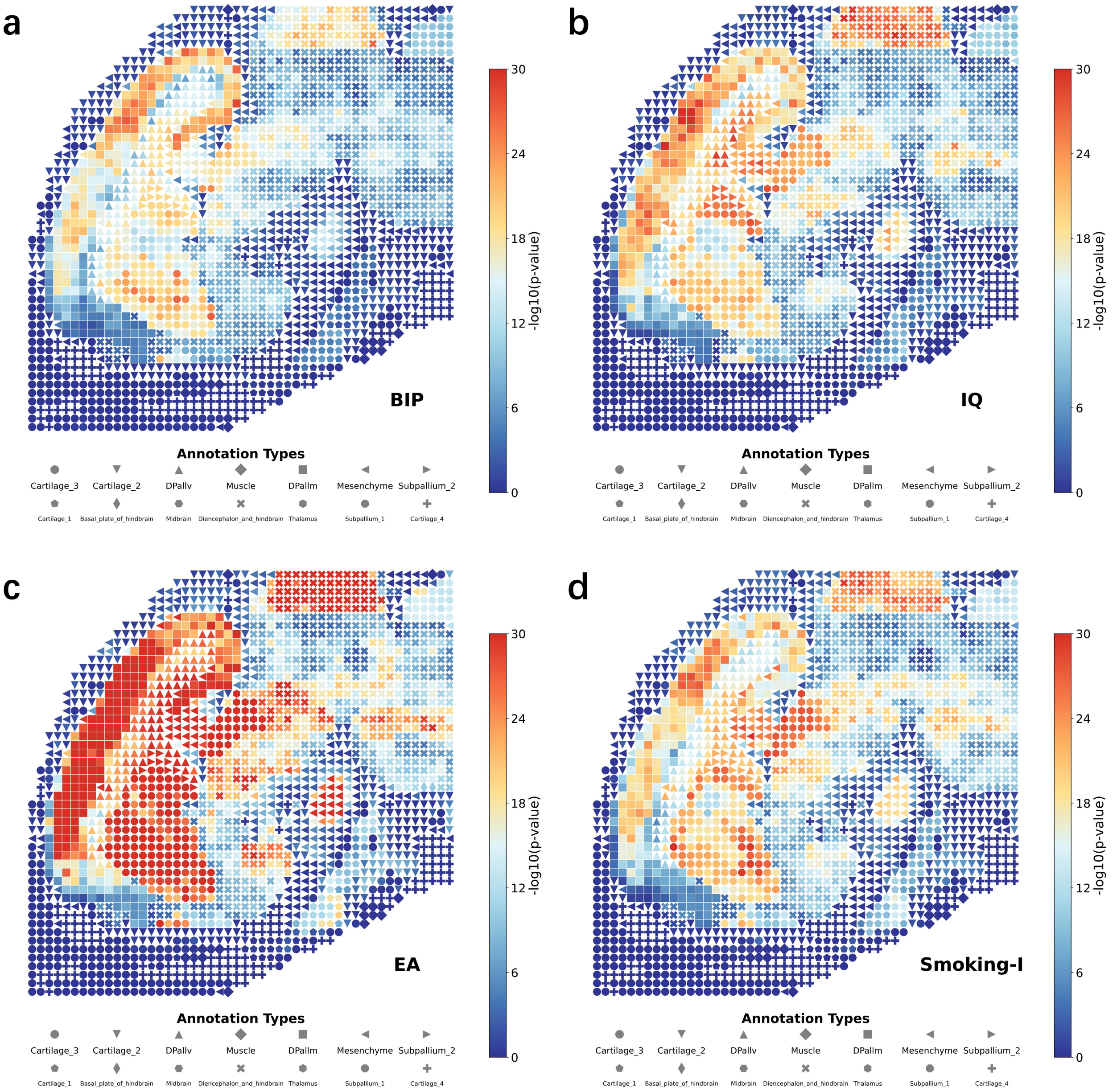


**Fig. S9**: Trait-spot associations for BIP, IQ, EA and Smoking-I traits identified by using gsMap embeddings in mouse brain datasets as input.

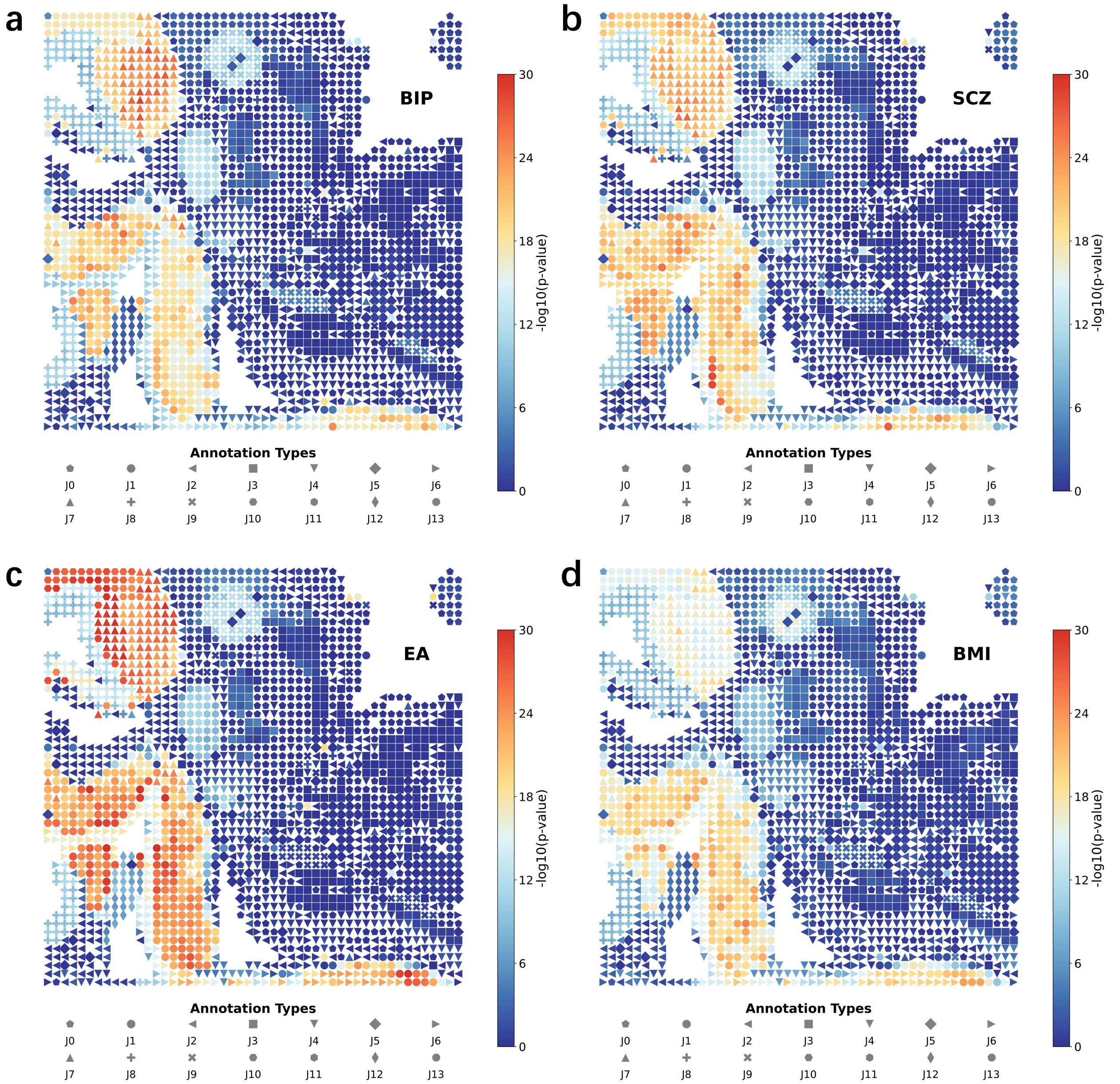


**Fig. S10**: Trait-spot associations for BIP, SCZ, EA, BMI traits identified by using CANDIES embeddings in mouse embryo datasets as input.

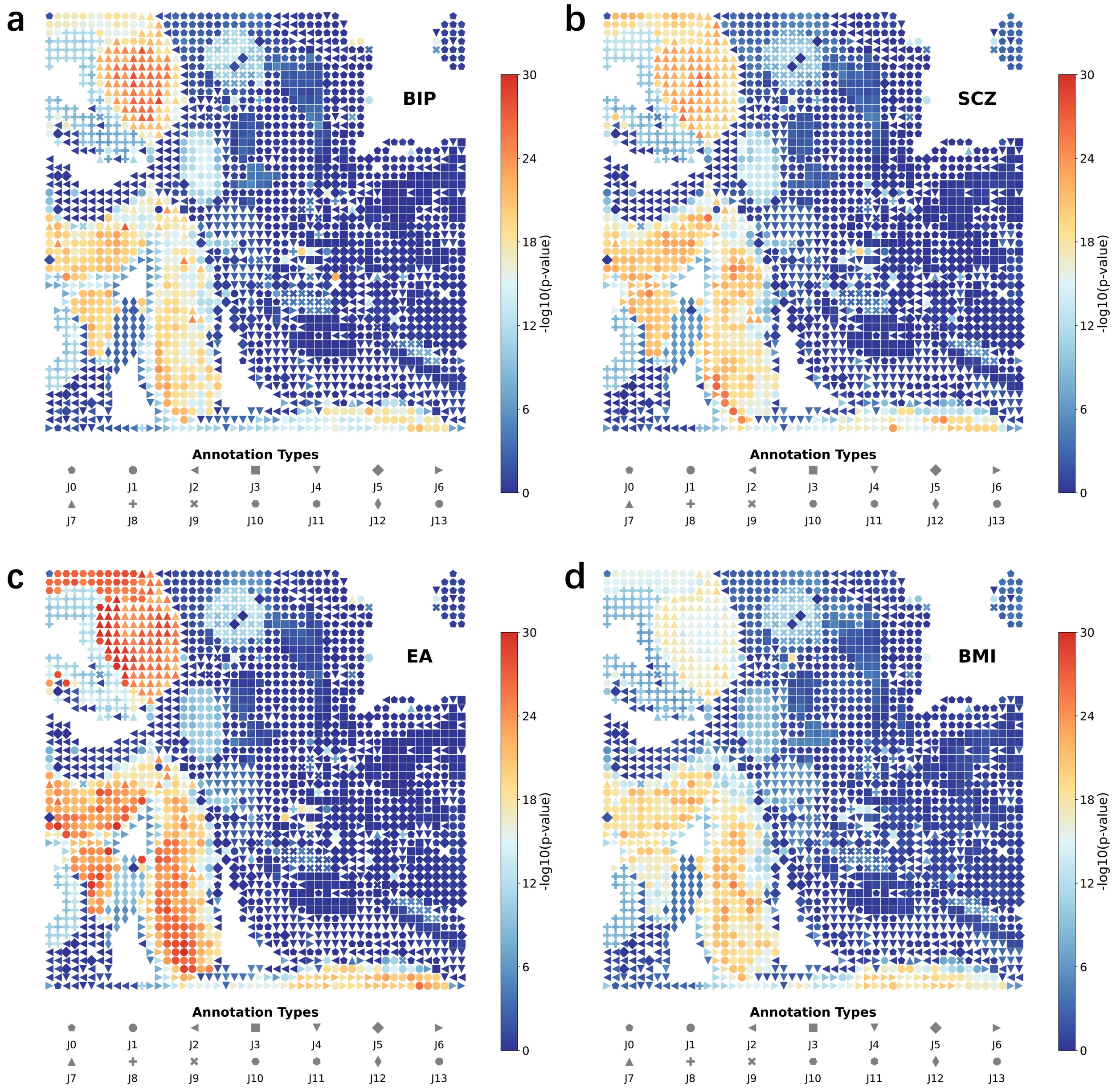


**Fig. S11**: Trait-spot associations for BIP, SCZ, EA, BMI traits identified by using gsMap embeddings in mouse embryo datasets as input.

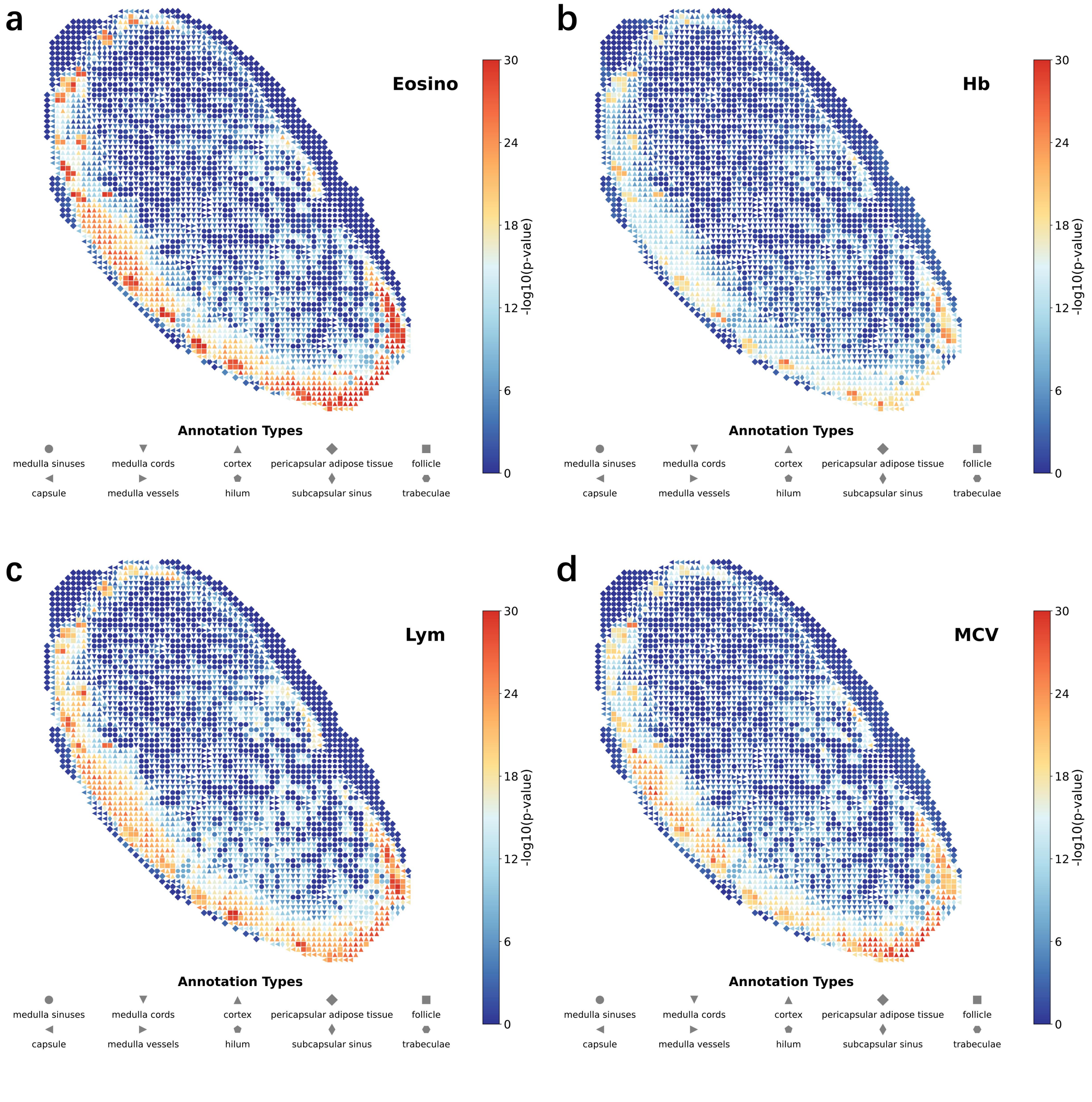


**Fig. S12**: Trait-spot associations for Eosino, Hb, Lym, MCV traits identified by using CANDIES embeddings in human lymph datasets as input.

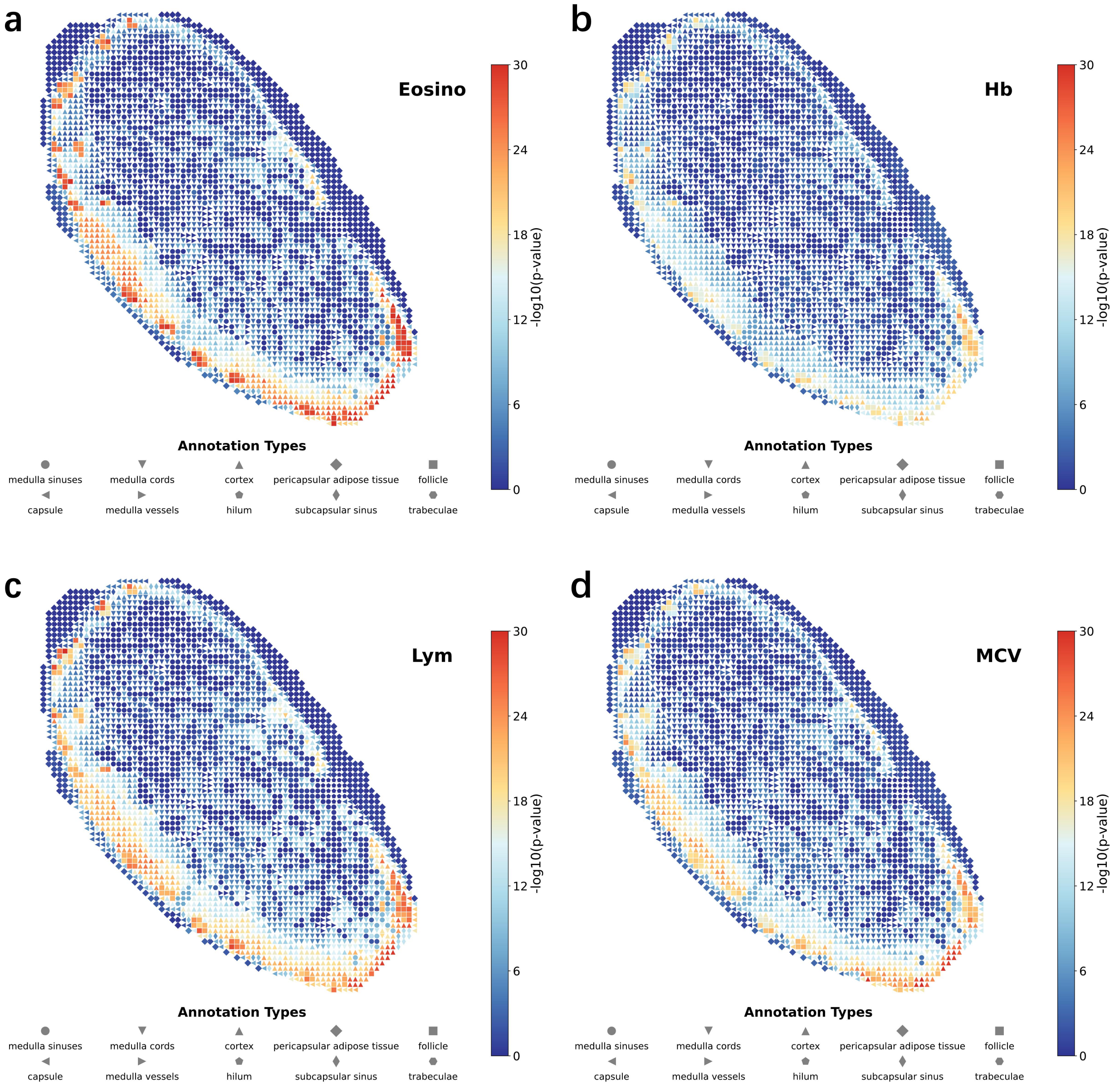


**Fig. S13**: Trait-spot associations for Eosino, Hb, Lym, MCV traits identified by using gsMap embeddings in human lymph datasets as input.

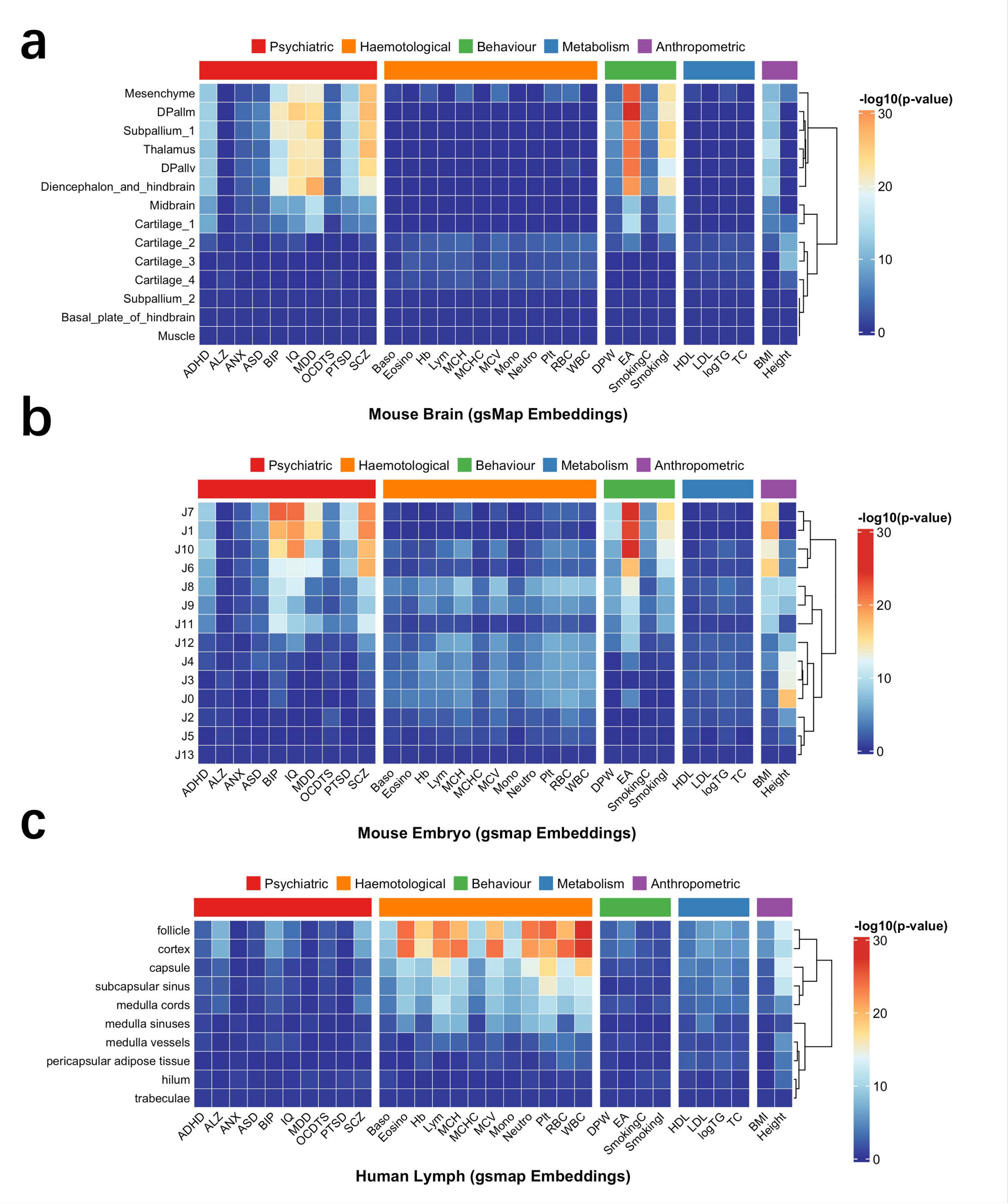


**Fig. S14**: Heatmaps showing the significance of associations between complex traits and spatial regions by using gsMap embedding.
